# Fat2 polarizes the WAVE complex in *trans* to align cell protrusions for collective migration

**DOI:** 10.1101/2022.01.17.476701

**Authors:** Audrey M. Williams, Seth Donoughe, Edwin Munro, Sally Horne-Badovinac

## Abstract

For a group of cells to migrate together, each cell must couple the polarity of its migratory machinery with that of the other cells in the cohort. Although collective cell migrations are common in animal development, little is known about how protrusions are coherently polarized among groups of migrating epithelial cells. We address this problem in the collective migration of the follicular epithelial cells in *Drosophila melanogaster*. In this epithelium, the cadherin Fat2 localizes to the trailing edge of each cell and promotes the formation of lamellipodia at the leading edge of the cell behind. We show that Fat2 performs this function by acting in *trans* to restrict WAVE complex activity to one long-lived region along each cell’s leading edge. Without Fat2, the WAVE complex distribution expands around the cell perimeter and fluctuates over time, resulting in reduced, unpolarized protrusive activity. We further show that Fat2’s influence is very local, with sub-micron-scale puncta of Fat2 concentrating the WAVE complex in corresponding puncta just across the leading-trailing cell-cell interface. These findings demonstrate that a trans interaction between Fat2 and the WAVE complex creates stable regions of protrusive activity in each cell and aligns the cells’ protrusions across the epithelium for directionally persistent collective migration.

## Introduction

Collective cell migration is essential for a variety of morphogenetic processes in animals^1–4^. As with individual cell migrations, adherent collective migrations are driven by the concerted action of cell protrusions, contractile actomyosin networks, and adhesions to a substrate^2,5,6^. To move forward, individual cells polarize these structures along a migratory axis, and to move persistently in one direction, they need to maintain that polarity stably over time^7^. Collective cell migrations introduce a new challenge: to move together, the group of migrating cells must be polarized in the same direction^7^. Otherwise, they would exert forces in different directions and move less efficiently, separate, or fail to migrate altogether.

The epithelial follicle cells of the *Drosophila melanogaster* ovary are a powerful experimental system in which to investigate how local interactions among migrating cells establish and maintain group polarity. Follicle cells are arranged in a continuous, topologically closed monolayer epithelium that forms the outer cell layer of the ellipsoidal egg chamber—the organ-like structure that gives rise to the egg^8^(Fig. 1). The apical surfaces of follicle cells adhere to a central germ cell cluster, and their basal surfaces face outward and adhere to a surrounding basement membrane extracellular matrix. The follicle cells migrate along this stationary basement membrane, resulting in rotation of the entire cell cluster^9^. As the cells migrate, they secrete additional basement membrane proteins^9^. The coordination of migration with secretion causes the cells to produce a basement membrane structure that channels tissue growth along one axis^9–12^. Follicle cell migration lasts for roughly two days, and the migration direction—and resulting direction of egg chamber rotation—is stable throughout^13,14^. The edgeless geometry of the epithelium means cells are not partitioned into “leader” and “follower” roles, and there is no open space, chemical gradient, or other external guidance cue to dictate the migration direction. Instead, this feat of stable cell polarization and directed migration is accomplished through local interactions between the migrating cells themselves^14,15^.

**Figure 1:**
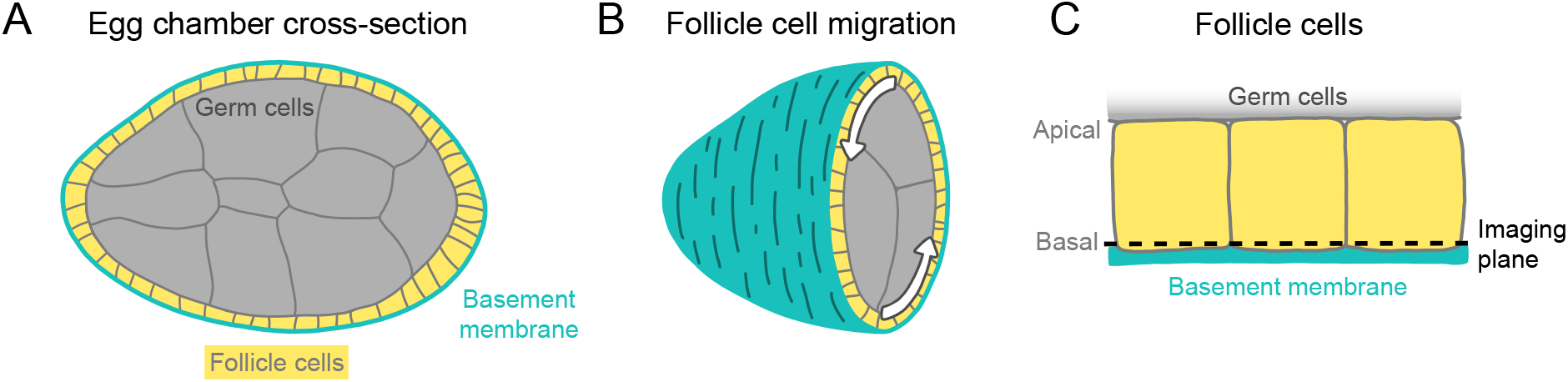
Introduction to egg chamber rotation. **A**, Diagram of a stage 6 egg chamber in cross-section. Anterior is left, posterior right. **B**, Three-dimensional diagram of an egg chamber with the anterior half shown. Arrows indicate the migration of follicle cells along the basement membrane and the resulting rotation of the egg chamber around its anterior-posterior axis. **C**, Diagram of three follicle cells. Their apical surfaces adhere to the germ cells and their basal surfaces adhere to the basement membrane. The dashed line represents the basal imaging plane used throughout this study except where indicated.

Follicle cell migration is driven, in part, by lamellipodial protrusions that extend from the leading edge of each cell^10,16^. Lamellipodia are built by the WASP family verprolin homolog regulatory complex (WAVE complex)^17,18^, which is a protein assembly composed of five subunits: SCAR/WAVE, Abi, Sra1/Cyfip, Hem/Nap1, and HSPC300^19^. The WAVE complex adds branches to actin filaments by activating the Actin-related proteins-2/3 complex (Arp2/3) and elongates existing filaments, building the branched actin network that pushes the leading edge forward^20–22^.

The follicle cells align their lamellipodial protrusions across the tissue, a form of planar polarity^10,16^. The atypical cadherin Fat2 is required both for this planar polarity and for collective migration to occur^23–25^. Fat2 is planar polarized to the trailing edge of each cell^24^, where it promotes the formation of protrusions at the leading edge of the cell immediately behind^15^. Interestingly, in addition to migration depending on polarized Fat2 activity, Fat2’s planar polarity also depends on epithelial migration^15^. It is not known how Fat2 regulates lamellipodia or cell polarity, or how these processes influence one another. We hypothesized that Fat2 acts as a coupler between tissue planar polarity and cell protrusion by polarizing WAVE complex activity to the leading edge of each cell. To test this, we used genetic mosaic analysis and quantitative imaging of fixed and live tissues to dissect Fat2’s contributions to protrusivity and protrusion polarity at cell and tissue scales.

We show that Fat2 signals in *trans*, entraining WAVE complex activity to one long-lived region along each cell’s leading edge. Without Fat2, the WAVE complex accumulates transiently at different regions around the cell perimeter, and cell protrusivity is reduced and unpolarized. The interaction between Fat2 and the WAVE complex is non-cell-autonomous but very local, with sub-micron-scale puncta of Fat2 along the trailing edge concentrating the WAVE complex just across the cell-cell interface, at the tips of filopodia embedded within the lamellipodium. These findings demonstrate how an intercellular interaction between Fat2 and the WAVE complex promotes cell protrusivity, stabilizes regions of protrusive activity along the cell perimeter, and aligns protrusions across the epithelium by coupling leading and trailing edges. Fat2-WAVE complex interaction thereby stabilizes the planar polarity of protrusions for directionally persistent collective migration.

## Results

### Fat2 increases and polarizes protrusions at the basal surface of the follicular epithelium

Recent work has shown that Fat2 regulates migration of the follicular epithelium by polarizing F-actin-rich protrusions; specifically, Fat2 at the trailing edge of each cell causes protrusions to form at the leading edge of the cell behind it, and without Fat2, protrusions are reduced or lost^15,26^. Beyond this qualitative description, it is not known how Fat2 modulates cell protrusion. Because protrusion is a dynamic process of spatially coordinated cell extension and retraction, we used live imaging and fluorescent labeling of the plasma membrane to obtain a more detailed, time-resolved view of protrusions and their distribution around cells. We acquired timelapse movies of the basal surface of control and *fat2N^103–2^* epithelia (a null allele, hereafter referred to as *fat2*) and saw that there was substantial protrusive activity in *fat2* epithelia, but the protrusions lacked the clear polarized distribution of control epithelia (Fig. 2A; Movie 1).

**Figure 2:**
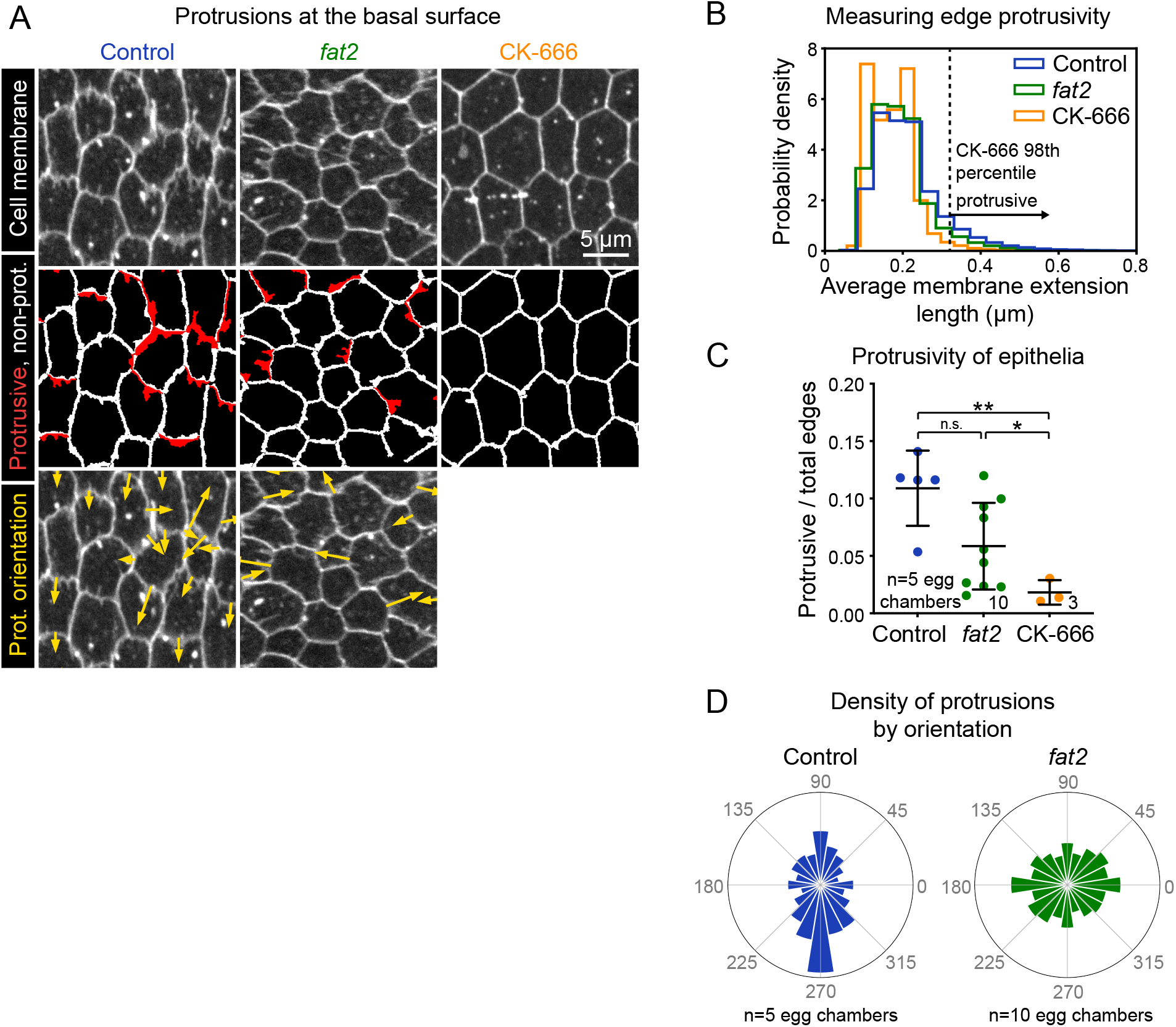
Fat2 increases and polarizes follicle cell protrusivity. **A**, Timelapse frames of control, *fat2*, and CK-666-treated epithelia labeled with a membrane dye. Middle row shows segmented edges. Protrusive edges, defined as edges with average membrane extension lengths longer than the 98th percentile of those of CK-666-treated epithelia, are shown in red. Non-protrusive edges are white. Bottom row shows arrows indicating the orientation of each protrusion overlaid on labeled cell membrane. Arrows originate at protrusion bases and have lengths proportional to protrusion lengths. See related Movies 1, 3. **B**, Histogram showing the distribution of average membrane extension lengths. The 98th percentile length threshold for CK-666-treated epithelia is indicated. **C**, Plot showing the ratio of protrusive to total edges. The protrusivity of *fat2* epithelia is variable, with a distribution overlapping with control and CK-666-treated epithelia. Welch’s ANOVA (W(2,8.5)=17, p=0.0011) with Dunnet’s T3 multiple comparisons test; n.s. p=0.07, *p=0.03, **p=0.006. **D**, Polar histograms of the distribution of protrusion orientations in control and *fat2* epithelia. Anterior is left, posterior is right, and in control epithelia images were flipped as needed so that migration is always oriented downward. Control protrusions point predominantly in the direction of migration, whereas *fat2* protrusions are less polarized. Associated with Figs. S1, S2, S3, S4; Movies 1, 3, 2, 4.

To quantify the extent and distribution of protrusions in these epithelia, we developed methods to segment membrane extensions and measure their lengths and orientations (Fig. S1). We briefly summarize our quantification approach here and include a more detailed explanation in the Methods and Materials. First, we measured the average lengths of membrane extensions from all cell-cell interfaces (Fig. 2B). The distribution of measured lengths was unimodal, with no natural division between protrusive and non-protrusive interfaces. Therefore, to establish an empirically-grounded cutoff between these categories, we recorded timelapse movies of control epithelia treated with the Arp2/3 inhibitor CK-666, which are non-migratory and almost entirely non-protrusive^16^. We used measurements from CK-666-treated epithelia to set a cutoff for the minimum length of a protrusion: any edges with membrane extensions longer than the 98th percentile of those in CK-666-treated epithelia were considered *protrusive* for subsequent analysis.

Using this quantification approach, we first asked how the amount of protrusions was affected by the loss of Fat2. We found that the protrusivity of *fat2* epithelia was lower than that of control on average, but highly variable, with overlap between the protrusivity distributions of both untreated and CK-666-treated epithelia (Fig. 2B,C; S2A,B; Movie 1). As a complementary method, we also measured protrusivity via F-actin labeling in fixed and live tissues, using *abi*-RNAi-express?ng epithelia as a nearly non-protrusive benchmark. The results largely paralleled those seen with membrane labeling (Fig. S3A-D; Movie 2); however, F-actin labeling showed a larger disparity in protrusivity between *fat2* and control epithelia (Figs. 2C; S3C). Images of follicle cell protrusions visualized by F-actin staining are dominated by fluorescence from filopodia, so the appearance of lower protrusivity of *fat2* epithelia as measured with an F-actin label may indicate that filopodia are disproportionately reduced by loss of Fat2. Altogether, these data show that *fat2* epithelia are less protrusive than control, but do retain some protrusive activity.

These results raised an important question—if some *fat2* epithelia have levels of membrane protrusivity comparable to that of control epithelia, then why do all *fat2* epithelia fail to migrate^13,15,24^? We hypothesized that the mispolarization of protrusions across the tissue contributes to *fat2* migration failure. In control epithelia, the majority of protrusions were polarized in the direction of migration, orthogonally to the egg chamber’s anterior-posterior axis (Fig. 2A,D; S2C; Movie 3). In contrast, in *fat2* epithelia, protrusions were fairly uniformly distributed in all directions or biased in two opposite directions (Fig. 2A,D; S2C; Movie 3). Where an axial bias was present, the axis was inconsistent between egg chambers. We also confirmed this finding using F-actin labeling of protrusions. To compare the planar polarity of F-actin protrusions between control and *fat2* epithelia, we measured F-actin enrichment at cell-cell interfaces as a function of the angle of the interface with respect to the egg chamber’s anterior-posterior axis. We again saw that protrusions were planar-polarized in control epithelia and unpolarized in *fat2* epithelia (Fig. S3A,E,F). These data show that Fat2 is required to polarize protrusions in a common direction across the epithelium.

Because Fat2 regulates both follicle cell migration and planar polarity, and migration and planar polarity are interdependent^15,16,23,24^, the unpolarized protrusions of *fat2* epithelia could be a cause or a consequence of inability of *fat2* epithelia to migrate. To distinguish between these possibilities, we exploited the fact that small groups of *fat2* cells can be carried along by neighboring non-mutant, migratory cells^24^, allowing us to evaluate polarity of protrusions from *fat2* cells in a migratory context. We generated *fat2* mosaic tissues that had sufficiently small fractions of mutant cells that the tissue as a whole still migrated, and found that *fat2* cells in these tissues were often protrusive, but their protrusions were not polarized in the direction of migration (Fig. S4; Movie 4). This demonstrates that Fat2 does not simply polarize protrusions indirectly by maintaining tissue-wide migration. Rather, Fat2 is required at the scale of groups of cells to polarize those cells’ protrusions in alignment with the direction of collective migration.

### Fat2 increases and polarizes the WAVE complex at the basal surface of the follicular epithelium

Follicle cell protrusions are built by the WAVE complex^16^, so we hypothesized that Fat2 polarizes protrusions by polarizing the WAVE complex’s distribution. To visualize the WAVE complex in living tissue and at normal expression levels, we used CRISPR/Cas9 to endogenously tag the WAVE complex subunit Sra1 with eGFP (hereafter: Sra1-GFP). We confirmed that Sra1-GFP flies are viable and fertile when the tagged allele is homozygous, Sra1-GFP localizes to follicle cell leading edges like other WAVE complex labels^16,26^, its localization depends on WAVE complex subunit Abi, and F-actin protrusions appear normal (Figs. 3A,B; S5A-C). Migration was slower when Sra1-GFP was present in two copies (Fig. S5D; Movie 5), so we performed all subsequent experiments with one copy of Sra1-GFP.

**Figure 3:**
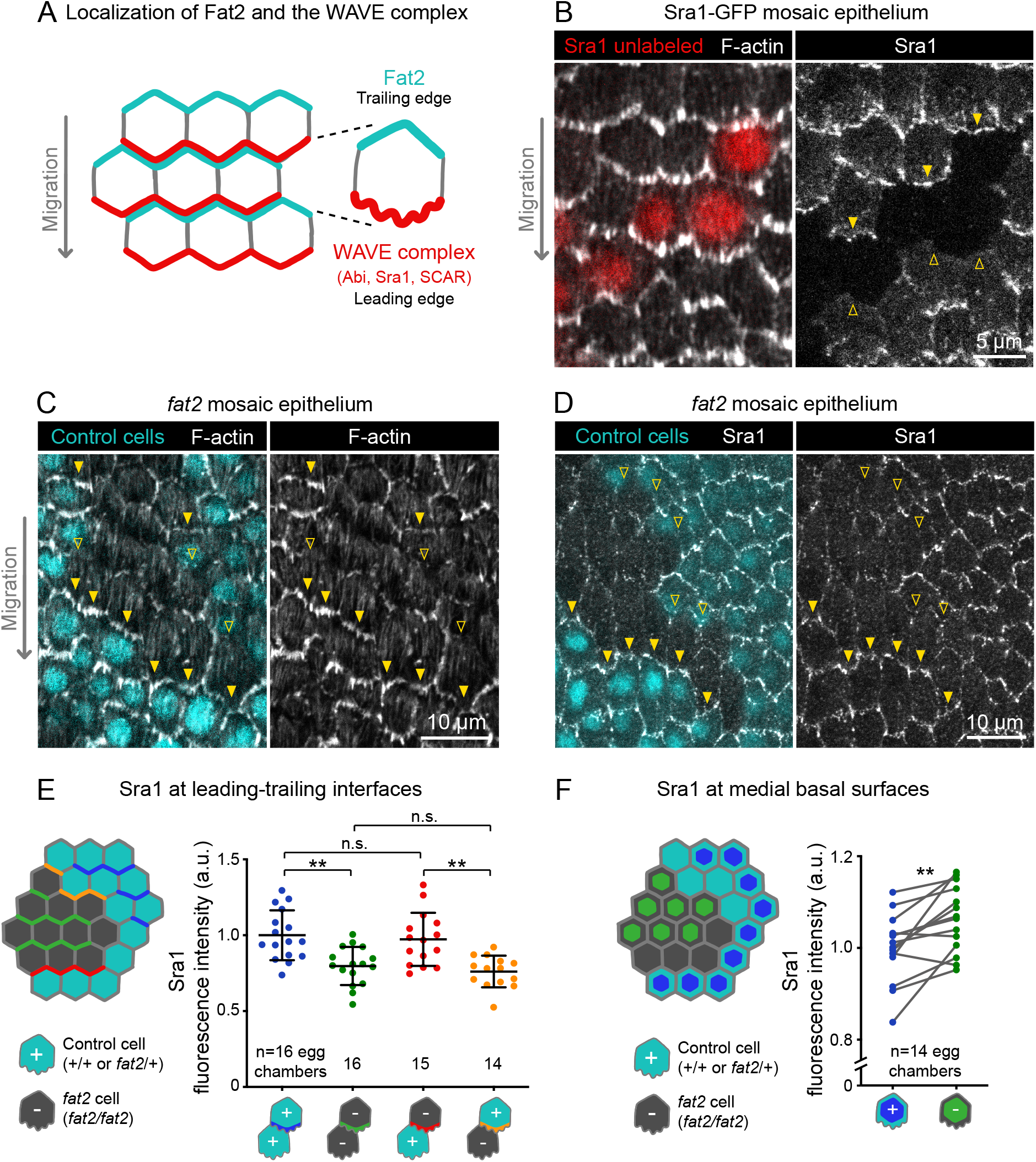
Fat2 in each cell concentrates the WAVE complex at the leading edge of the cell behind. **A**, Diagram showing Fat2 localization at the trailing edge and WAVE complex at the leading edge of the basal surface of follicle cells. The WAVE complex subunits referenced in this paper listed. **B**, Images of an Sra1-GFP mosaic epithelium with phalloidin-stained Factin, showing Sra1-GFP enrichment at leading edges (filled arrows) and not trailing edges (open arrows). **C**, Images of a *fat2* mosaic epithelium with phalloidin-stained F-actin. Filled arrows indicate leading edges of *fat2* cells behind control cells, where protrusions are present. Open arrows indicate leading edges of control cells *behind fat2* cells, where protrusions are reduced. **D**, Images of *afat2* mosaic epithelium expressing Sra1-GFP. Filled arrows indicate leading edges of *fat2* cells behind control cells. Open arrows indicate leading edges of control cells behind *fat2* cells. **E,F**, Quantification of Sra1-GFP mean fluorescence intensity in *fat2* mosaic epithelia along leading-trailing interfaces (E) or medial basal surfaces (F). Diagrams to the left of plots show the measured regions with respect to control (cyan) *and fat2* (gray) cells. The genotype(s) of cells in each measured category are shown below the x-axis. **E**, Sra1-GFP is reduced at the leading edge of cells of any genotype behind *fat2* cells. Bars indicate mean ±SD. One-way ANOVA (F(3,57)=10.40, p<0.0001) with post-hoc Tukey’s test; n.s. (left to right) p=0.96, 0.90, **p<0.01. **F**, Sra1-GFP is slightly increased at the medial basal surface of *fat2* cells. Lines connect measurements from the same egg chamber. Paired t-test; **p<0.01. Associated with Figs. S5, S6; Movie 5.

With an endogenous WAVE complex label in hand, we investigated how Fat2 affects WAVE complex localization. Previous work has shown that WAVE complex levels are reduced at the basal surface of follicle cells lacking Fat2^26^. Consistent with this result, we found that Sra1-GFP levels were lower along cell-cell interfaces at the basal surface of *fat2* epithelia than of control epithelia (Fig. S6A-C). Planar polarity of Sra1-GFP across the epithelium was also lost in the absence of Fat2 (Fig. S6D,E). Fat2 acts non-cell-autonomously to cause protrusions to form at the leading edge of the cell just behind^15^ (Fig. 3C), so we next tested the hypothesis that Fat2 localizes the WAVE complex to the leading edge in the same non-cell-autonomous pattern. We did this using *fat2* mosaic epithelia, in which we could measure Sra1-GFP levels at leading-trailing interfaces shared by control and *fat2* cells. We found that Sra1-GFP levels were normally-enriched along the leading edges of *fat2* cells if control cells were present immediately ahead, showing that Sra1 can still localize to the leading edge of cells lacking Fat2. Conversely, Sra1-GFP levels were reduced along the leading edge of control cells if *fat2* cells were immediately ahead (Fig. 3D,E). We also observed a corresponding non-autonomous pattern of membrane protrusion polarity in timelapse movies of *fat2* mosaic epithelia (Fig. S4; Movie 4). We conclude that Fat2 acts non-cell-autonomously to localize the WAVE complex to leading edges, resulting in tissue-wide planar polarization of protrusive activity, and thereby in collective cell migration.

We next asked if loss of the WAVE complex from the leading edge in the absence of Fat2 meant that the protein was redistributed to other cell surfaces. To test this, we compared the level of Sra1-GFP at the medial basal surfaces of control and *fat2* cells in mosaic epithelia (see diagram in Fig. 3F). We found that Sra1-GFP levels were slightly increased in *fat2* cells compared to control cells in the same tissue (Fig. 3D,F). There was not a statistically significant increase in Sra1-GFP levels at medial basal cell surfaces of epithelia composed entirely of control or *fat2* cells, although the trend was the same (Fig. S6C). These data demonstrate that Fat2 concentrates the WAVE complex at the leading edge, and that without Fat2, the WAVE complex becomes distributed more broadly across the basal surface.

### Fat2 stabilizes a region of WAVE complex enrichment and protrusivity in *trans*

In individually migrating cells, the excitable dynamics of the WAVE complex and its regulators enable it to form transient zones of enrichment along the cell perimeter even in the absence of a directional signal^7,27,28^. Although the planar-polarized distribution of the WAVE complex across the epithelium was lost in *fat2* mutant tissue, we wondered (1) whether the WAVE complex could still form regions of enrichment in individual cells and (2) whether these WAVE complex-enriched regions were active and responsible for templating unpolarized protrusions. To evaluate the WAVE complex distribution along the edges of individual cells, we generated entirely *fat2* mutant epithelia in which patches of cells expressed Sra1-GFP. At cell-cell interfaces along Sra1-GFP expression boundaries, we found that the boundary cells often had cortical regions devoid of Sra-GFP (Fig 4A). This observation shows that the WAVE complex is not uniformly localized around the cortex and can form regions of enrichment without Fat2. We also saw that Sra1-GFP enrichment coincided with the presence of F-actin protrusions (Fig. 4A), indicating that the WAVE complex in these regions is active. To confirm that the WAVE complex builds the protrusions in *fat2* epithelia, we co-imaged Sra1-GFP and a membrane label, and found that Sra1-GFP was enriched at the tips of membrane protrusions (Fig. S7A; Movie 6). These data indicate that the WAVE complex can still accumulate and build protrusions in the absence of Fat2, tissue-wide planar polarity, and collective cell migration.

**Figure 4:**
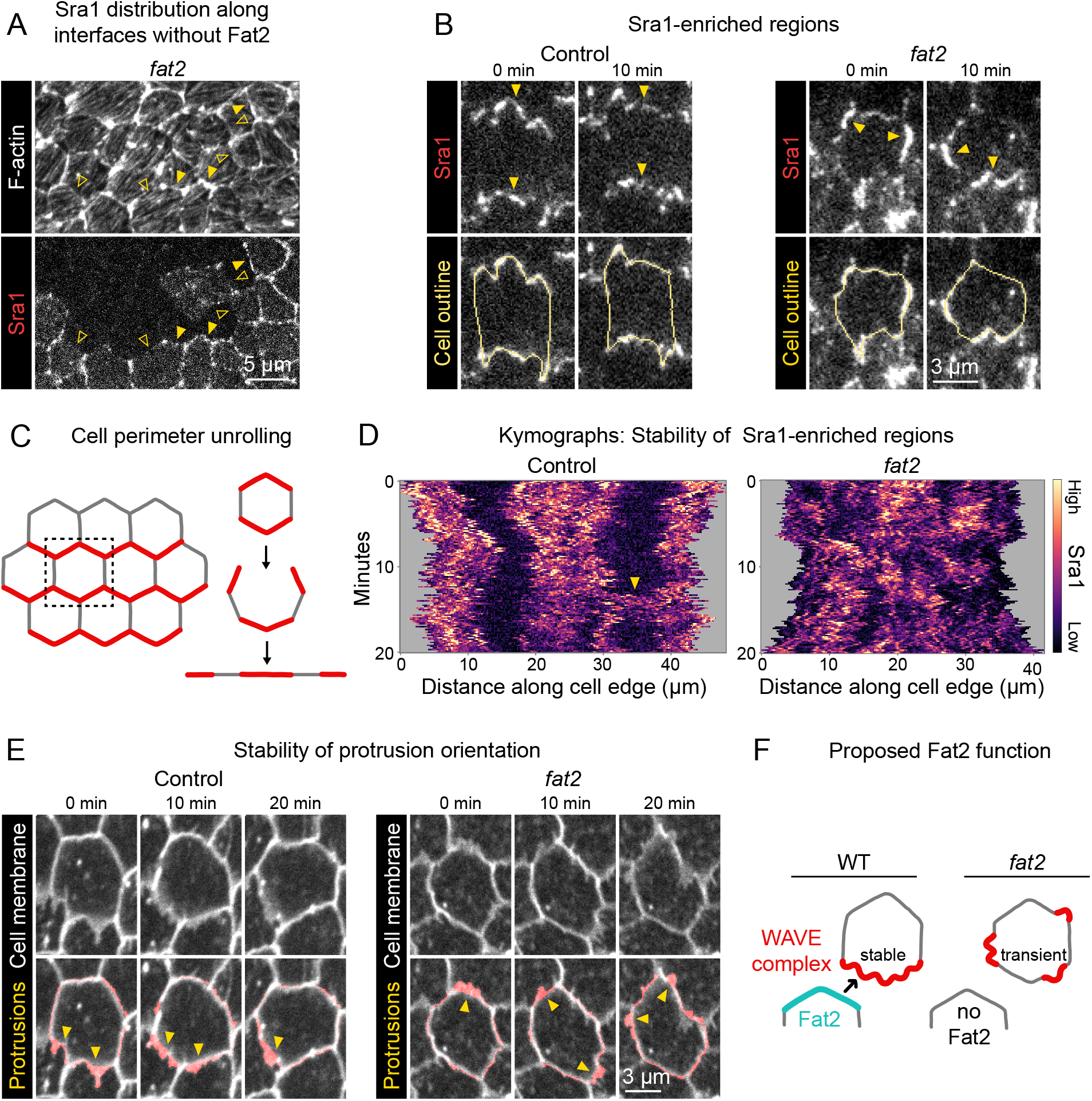
Fat2 stabilizes a region of WAVE complex enrichment and protrusivity. **A**, Images of phalloidin-stained Factin and mosaically-expressed Sra1-GFP in an entirely *fat2* mutant epithelium. Filled and open arrows indicate genotype boundary interfaces with and without Sra1-GFP enrichment, respectively. Sra1-GFP enrichment is heterogeneous, and interfaces with Sra1-GFP enrichment have more F-actin protrusions. **B**, Timelapse frames of Sra1-GFP in control and *fat2* epithelia. Top row shows Sra1-GFP with arrows indicating regions of Sra1-GFP enrichment; bottom row shows Sra1-GFP and outlines of cell perimeters used to make kymographs. Laser intensity and brightness display settings differ between genotypes. See related Movies 7, 8.**C**, Diagram of cell perimeter unrolling for kymograph generation. Red represents planar-polarized Sra1 as distributed before and after unrolling. **D**, Kymographs of Sra1-GFP fluorescence intensity along cell perimeter outlines exemplified in (C). The *y*-axis length of regions of high Sra1-GFP enrichment reports their stability over time. Control cells have Sra1-GFP regions along leading-trailing interfaces that are stable over 20 minutes. In *fat2* cells, Sra1-GFP-enriched regions are less stable. The arrow indicates a transient accumulation of Sra1-GFP at a control cell side. These occur occasionally, and their stability is similar to Sra1-GFP regions in *fat2* cells. **E**, Timelapse frames of control and *fat2* epithelia with a membrane dye. Top row shows the interfaces and protrusions of one cell and its neighbors. Segmented membrane extensions originating from the center cell (red) are overlaid in the bottom row. Arrows indicate sites of membrane protrusion. The position of protrusions in the *fat2* cell changes more than in the control cell. See related Movie 10. **F**, Diagram showing the proposed role of Fat2 stabilizing a region of WAVE complex enrichment and protrusivity. Without Fat2, WAVE complex-enriched, protrusive regions are reduced and more transient. Associated with Fig. S7; Movies 6, 7, 8, 10, 9.

A striking feature of migrating follicle cells is the stable polarization of their protrusive leading edges. It is not known whether Fat2 contributes to the stabilization of protrusive regions in addition to positioning them. If so, the positions of protrusive regions of *fat2* epithelia should fluctuate more than those of control epithelia, in addition to being less well-polarized at the tissue level. To see if this is the case, we acquired timelapse movies of Sra1-GFP and monitored its distribution along cell perimeters over time. In control epithelia, Sra1-GFP was strongly enriched along leading-trailing interfaces relative to side interfaces over the 20-minute timelapse. Side interfaces were mostly devoid of Sra1-GFP, except for infrequent Sra1-GFP accumulations that persisted for several minutes (Fig. 4B-D; Movies 7; 8). In contrast, in *fat2* epithelia, the regions of greatest Sra1-GFP enrichment along the cell perimeter changed substantially over the 20-minute timelapse and multiple Sra1-GFP-enriched regions were often present simultaneously in individual *fat2* cells. Sra1-GFP accumulated in these regions, typically spreading outward along the membrane as it did so, and then dissipated. These events had a duration that was comparable to the transient accumulations of Sra1-GFP at side interfaces in control cells (Fig. 4B-D; Movies 7;8). Because all cell-cell interfaces in *fat2* epithelia and side interfaces in control epithelia lack Fat2, this suggests a several-minutes timescale over which regions of WAVE complex enrichment can persist without stabilization by Fat2. Live imaging of Sra1-GFP in *fat2* mosaic epithelia yielded similar information—Sra1-GFP enrichment fluctuated more at interfaces between *fat2* cells than interfaces between control cells despite both being in a migratory tissue (Fig. S7B,C; Movie 9). To see if Fat2’s role stabilizing the WAVE complex distribution translates to a role stabilizing protrusive regions, we compared the stability of regions of membrane protrusion between cells in control or *fat2* epithelia. In control cells, we found that protrusions extended and retracted from one leading edge region, and changes to the direction of cell protrusion were rare (Fig. 4E; Movie 10). In contrast, the edges of cells in *fat2* epithelia undergoing protrusion often shifted substantially over the 20 minutes of the timelapse. Together, these observations show that, in addition to polarizing protrusive activity to the leading edge, Fat2 stabilizes the distribution of WAVE complex activity for repeated cycles of protrusion from one long-lived protrusive region (Fig 4F).

### Fat2 and the WAVE complex colocalize across leading-trailing cell-cell interfaces

Finally, we explored how Fat2 recruits the WAVE complex across the cell-cell interface. To constrain the set of possible mechanisms, we assessed the spatial scale of their interaction. Fat2 has a punctate distribution along each cell’s trailing edge^15,24^, so we asked whether Fat2 recruits the WAVE complex locally to these sites, or recruits it more broadly to the entire interface. We evaluated the colocalization between Fat2 and the WAVE complex along leading-trailing interfaces, visualizing Fat2 with an endogenous 3xeGFP tag (Fat2-3xGFP) and the WAVE complex with mCherry-tagged Abi under control of the ubiquitin promoter (Abi-mCherry). Like Fat2-3xGFP, Abi-mCherry formed puncta, and Abi-mCherry and Fat2-3xGFP puncta colocalized significantly more than Abi-mCherry and uniformly-distributed E-cadherin-GFP (Spearman’s r = 0.71±0.04 vs. 0.49±0.07; Figs 5A-E; S8A,B). In timelapse movies, Fat2-3xGFP and Abi-mCherry puncta moved together through cycles of protrusion extension and retraction (Figs. 5B; Movie 11). Short-lived Abi-mCherry accumulations formed infrequently at cell sides away from Fat2, similar to the Sra1-GFP side accumulations we described earlier (Figs. 4D; S9; Movies 7; 12). Together, these findings suggest that Fat2 recruits the WAVE complex locally, at the scale of individual puncta, with the WAVE complex occasionally “escaping” Fat2-dependent concentration at the leading edge.

If Fat2 puncta locally recruit the WAVE complex, changing the distribution of Fat2 puncta should cause corresponding changes to the distribution of the WAVE complex. To test this, we examined follicle cells expressing an endogenous Fat2 truncation that lacks the intracellular domain (Fat2^ΔICD^-3xGFP), which distributes more broadly around the cell perimeter than wild-type Fat^15,29^, but remains punctate. The distribution of Abi-mCherry expanded around the cell perimeter in the Fat2^ΔICD^-3xGFP background (Fig. 5A) as was previously reported for protrusions^15^. Despite their altered distributions, Abi-mCherry puncta colocalized just as well with Fat2^ΔICD^-3xGFP puncta as with Fat2-3xGFP puncta (Spearman’s r = 0.71±0.04 vs. 0.71±0.05; Figs. 5E; S8B). From these data we conclude that Fat2 controls the distribution of the WAVE complex by concentrating the WAVE complex in adjacent puncta. These findings also demonstrate that the Fat2 intracellular domain is dispensable for Fat2-WAVE complex interaction in collectively migrating follicle cells.

**Figure 5:**
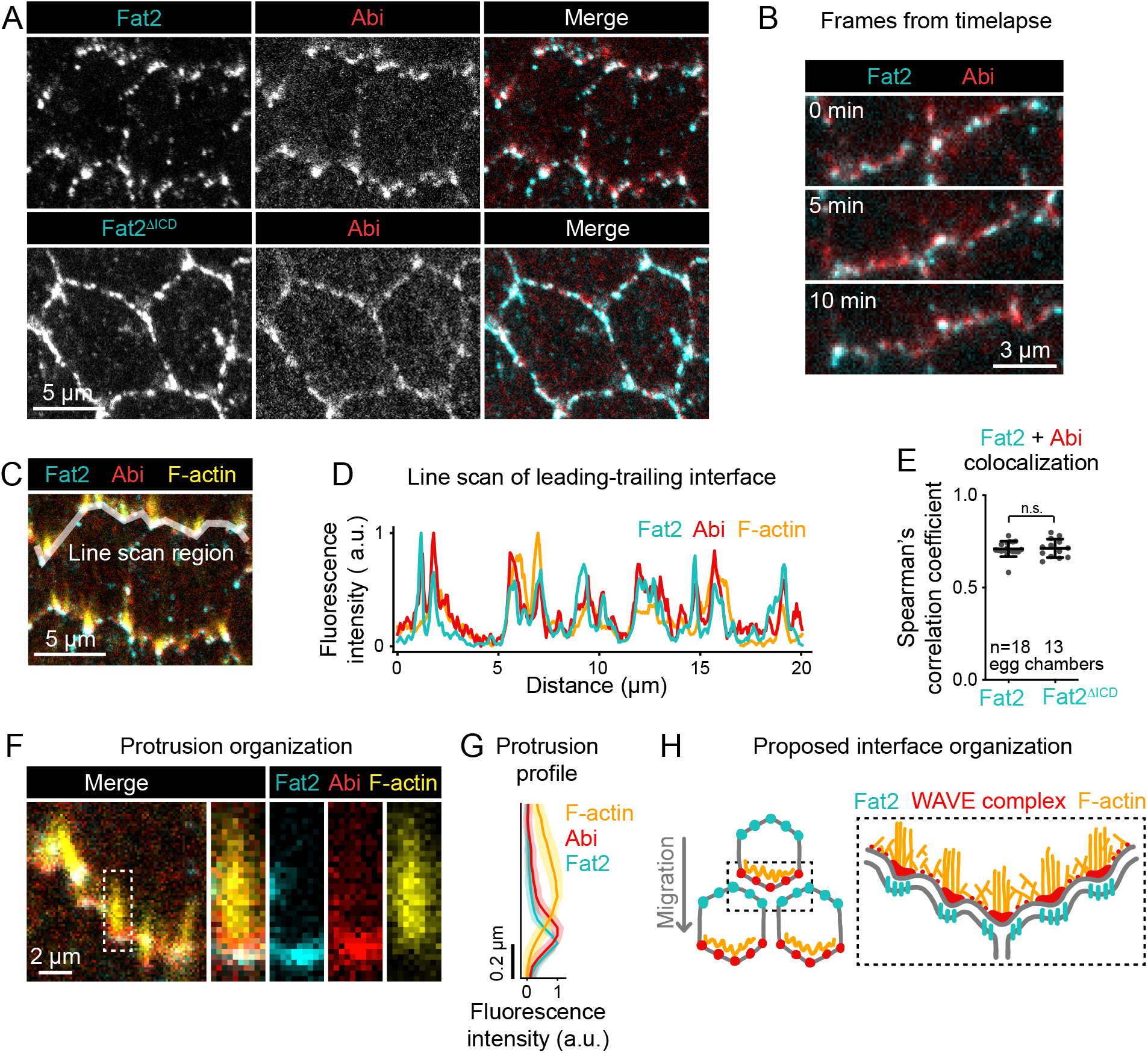
Fat2 colocalizes with the WAVE complex across leading-trailing cell-cell interfaces. A, Images of cells expressing Abi-mCherry and endogenous full-length Fat2-3xGFP or endogenous Fat2-3xGFP lacking the intracellular domain (Fat2^ΔICD^), used to assess colocalization. B, Timelapse frames showing the leading-trailing interfaces of two cells expressing Fat2-3xGFP and Abi-mCherry, showing their colocalization over time. See related Movie 11. C, Image showing the leading-trailing interface region used in (D); it is also an example of a region used in (E). D, Line scan showing the fluorescence intensity of Fat2-3xGFP, Abi-mCherry, and F-actin (phalloidin) along the leading-trailing interfaces of the two cells in (C), showing their corresponding peaks of enrichment. E, Plot of Spearman’s correlation coefficients of Fat2-3xGFP or Fat2^ΔICD^-3xGFP and Abi-mCherry showing no significant difference in colocalization. Bars indicate mean ± SD. Oneway ANOVA (F(5,81)=44.86, p=0.0164 with Fig. S8B) with post-hoc Tukey’s test; n.s. p>0.99. F, Image showing the distribution of Fat2-3xGFP, Abi-mCherry, and F-actin (phalloidin) at the leading-trailing interface and along the boxed filopodium. G, Plot showing fluorescence intensity of traces of F-actin, Abi-mCherry, and Fat2-3xGFP showing their relative sites of enrichment along the length of filopodia. Lines and shaded regions indicate mean ± SD. n=74 protrusions (used for SD), 18 cells, 1 cell/egg chamber. H, Diagram of proposed organization of Fat2, the WAVE complex, and F-actin along the leading-trailing interface based on the present data and previously published work^15,16,24^. Fat2 puncta at the trailing edge colocalize with WAVE complex puncta at the leading edge, ahead of filopodia embedded within the lamellipodium. Associated with Figs. S8, S9; Movies 11, 12.

Ena-dependent filopodia are embedded within and grow from the lamellipodia^16^. The WAVE complex interacts with Ena and is required for the filopodia to form^16,30^, so we asked whether the distribution of Fat2-WAVE complex puncta is related to the distribution of filopodia. Labeling filopodia tips with a GFP-tagged Ena transgene (GFP-Ena) and comparing the localization of Abi-mCherry and F-actin to either GFP-Ena or to Fat2-3xGFP, we found that the sites of highest Fat2-3xGFP and Abi-mCherry enrichment coincided with filopodia tips (Figs. 5C,D,F; S8C,D). Fluorescence intensity profiles along filopodia lengths showed that Fat2-3xGFP and Abi-mCherry were enriched just ahead of the F-actin-rich region (Fig. 5F,G). Fat2-3xGFP was shifted slightly forward from Abi-mCherry, consistent with the separation of Fat2-3xGFP and Abi-mCherry fluorophores by a cell-cell interface (Figs. 5F,G; S8D,E). This analysis demonstrates a stereotyped organization in which Fat2 and the WAVE complex are concentrated along with Ena near the tips of the filopodia, with Fat2 at the trailing edge across the cell-cell interface from the leading edge components.

We considered two explanations for the close spatial relationship between Fat2 puncta, WAVE complex puncta, and filopodia. Fat2 could recruit the WAVE complex locally to puncta, and WAVE complex puncta shape the distribution of filopodia. Alternatively, Fat2 could recruit the WAVE complex to the leading edge, but their colocalization in puncta be a secondary effect of the filopodia, perhaps caused by the known interaction between Ena and Abi^30^ or by deformation of the leading-trailing interface. To rule out the second possibility, we measured colocalization between Fat2-3xGFP and Abi-mCherry in *ena*-RNAi-expressing epithelia, in which filopodia are strongly depleted^16^ (Fig. S8F). Despite the loss of filopodia, both Fat2-3xGFP and Abi-mCherry remained punctate, and their colocalization was only slightly reduced (Spearman’s r = 0.71±0.04 vs. 0.65±0.03, Fig. S8A,B). We therefore rule out Ena or the filopodia themselves as required mediators of the spatial relationship between Fat2 and the WAVE complex, and infer that Fat2-WAVE complex colocalization is indicative of Fat2 recruitment of the WAVE complex locally to these sites.

Altogether, we propose that Fat2 acts locally, at the scale of individual Fat2 puncta, to concentrate the WAVE complex in adjacent puncta across the cell-cell interface. Because Fat2 puncta are distributed along the trailing edge, this has the broader effect of stabilizing a region of WAVE complex enrichment at the leading edge.

## Discussion

This work demonstrates that a *trans* interaction between the atypical cadherin Fat2 and the WAVE complex can stabilize WAVE complex polarity for directed cell migration. Fat2, localized to the trailing edge of each cell, recruits the WAVE complex to the leading edge of the cell behind, just across their shared interface. By concentrating WAVE complex activity in a restricted region, Fat2 strongly biases lamellipodia and filopodia to form at these leading edge sites, stably polarizing overall cell protrusive activity to one cell side. Because the Fat2-WAVE complex signaling system is deployed at each leading-trailing interface in a planar-polarized manner, it both polarizes protrusions within individual cells and aligns these individual cell polarities across the epithelium. This allows the cells to exert force in a common direction and achieve a highly coordinated collective cell migration.

While the molecular players differ, local coupling of leading and trailing edges through asymmetric interactions across their shared interface is a recurring motif in studies of epithelial collective cell migrations. In an epithelial cell culture model of collective migration, asymmetric pulling forces across cell-cell interfaces polarize Rac1 activity and cell protrusion^31^. In another model, one cell’s lamellipodium is stabilized by confinement under the trailing edge of the cell ahead, reinforcing interface asymmetry^32^. In an endothelial collective cell migration model, asymmetric membrane “fingers” containing VE-cadherin extend from the trailing edge and are engulfed by the leading edge of the cell behind, whose movement they help guide^33^. These types of leading-trailing edge coupling systems could operate together with longer-range cues to reinforce the planar polarity of cells’ migratory structures. In migrations with a closed topology and no extrinsic directional cues, such as that of the follicle cells, local polarity coupling may be especially critical for collective migration.

Our development of new computational tools to segment and quantify membrane protrusion dynamics in a collectively-migrating epithelium has led to new insights into how Fat2 regulates protrusions. We found that without Fat2, protrusivity was reduced, and the distribution of remaining protrusions expanded around the entire cell periphery. Therefore, Fat2 not only promotes protrusion at the leading edge, but also restricts protrusion to that edge. Analysis of *fat2* mosaic epithelia revealed that Fat2 acts locally to enforce this restriction—even in the context of a globally planar-polarized, migratory epithelium, cells lacking input from Fat2 in the cell ahead were unable to polarize their protrusions in the direction of migration. Although we focused on the length and orientation of protrusions in this study, our approach could be extended to other protrusion traits such as shape, lifetime, or elongation rate, and/or to other tissues, making it broadly applicable to studies of epithelial collective migrations.

Excitable WAVE complex dynamics underlie lamellipodial protrusion^27,34,35^, and these dynamics were especially apparent in our data in contexts where Fat2 was absent from the cell-cell interface—either at interfaces between cells in *fat2* epithelia, or at protrusive side interfaces we observed with low frequency in control cells. In both contexts, the WAVE complex accumulated at an edge region, spread laterally along the membrane, and then dissipated. This corresponded with the initiation, growth, and collapse of a protrusion. Where Fat2 was present, the WAVE complex distribution along the cell perimeter stayed more constant and WAVE complex levels fluctuated in place, but did not appear to spread laterally. In the absence of Fat2, the WAVE complex also had an expanded distribution around lateral cell edges and across the basal surface. These findings suggests that Fat2 acts by sequestering the WAVE complex, reducing its ability to accumulate elsewhere and thus restricting protrusions to a single leading edge. Moreover, the reduced protrusivity of *fat2* epithelia may be due to their broader and more diffuse WAVE complex distribution, which could reduce the frequency with which the WAVE complex crosses a threshold of enrichment necessary to initiate a protrusion. The sporadic side-facing protrusions in wild-type cells show that the Fat2 signaling system does not exert perfect control over the distribution of WAVE complex activity, but this level of control is sufficient to stably polarize cell protrusive activity for days-long, highly directed migration.

Given that follicle cells can form WAVE complex-dependent protrusions without Fat2, how does Fat2 shape the WAVE complex’s distribution and dynamics in wild-type cells? WAVE complex activity is often entrained by directional cues from the environment^34,36–38^, and we hypothesize that Fat2 is acting as a similar directional cue in follicle cells. The WAVE complex is activated by recruitment to the plasma membrane^19,39,40^. Positive regulators of WAVE complex accumulation include active Rac, phosphatidylinositol (3,4,5)-triphosphate (PIP_3_), membrane-localized proteins that directly bind the WAVE complex, and the WAVE complex itself^17,35,39,41–46^. We propose that Fat2 promotes WAVE complex accumulation within a stable region by acting through one or more of these positive regulators, thereby controlling the site where the WAVE complex excitation threshold is crossed and a protrusion is formed. Under this model, in the absence of Fat2, this site selection instead becomes more stochastic and therefore long-lasting protrusive regions cannot form.

Fat2 acts at the trailing edge of each cell to recruit the WAVE complex in *trans*, so there must be one or more transmembrane proteins at the leading edge of each cell that bridge this interaction. Previous work has shown that the receptor tyrosine phosphatase Lar is part of this bridge—Fat2 recruits Lar to each follicle cell’s leading edge^15^, and in Lar’s absence both WAVE complex levels and cell protrusions are reduced^15,26^ (Fig. S8G). However, the WAVE complex that persists at the leading edges of *lar* cells still colocalizes with Fat2 (Fig. S8A,B). Therefore, there must be at least one other transmembrane protein that works alongside Lar to mediate the Fat2-WAVE complex interaction. Identifying the missing leading edge protein(s) will be important to fully understand how Fat2 shapes WAVE complex activity.

Fat2 is localized in puncta along each cell’s trailing edge^15,24^, and we show here that these puncta correspond 1:1 with regions of high WAVE complex enrichment just across the leading-trailing cell-cell interface. Fat2’s punctate distribution and its levels along cell-cell interfaces are unaffected by loss of the WAVE complex^15^, indicating that Fat2 puncta shape the distribution of the WAVE complex and protrusions, not the reverse. We further show that the puncta sit at the tips of filopodia that form within the lamellipodial actin network. Filopodia are a prominent feature of the long-lived protrusive regions that form in wild-type epithelia, but appear to be disproportionately reduced in the short-lived, fluctuating protrusive regions that form in fat2 epithelia. We therefore propose that by concentrating the WAVE complex and/or stabilizing its distribution, Fat2 also facilitates filopodia formation. It should be noted, however, that the filopodia are dispensable for collective follicle cell migration^16^, so the reason these structures form remains to be determined.

Why, and how, is Fat2 localized in puncta? Cadherins commonly form puncta, though the causes and functions of this organization vary^47–49^. For example, Flamingo (or mammalian Celsr1), an atypical cadherin and central component of the core planar cell polarity pathway, is stabilized by clustering, and this clustering is important for its planar polarization^50–52^. In future work, it will be important to determine how Fat2 assembles in puncta, and whether this local clustering is important for its polarization to trailing edges or its effect on the organization of leading edges. More broadly, it will be critical to determine how Fat2 achieves its trailing edge localization, a necessary step in the polarization of the tissue.

## Author Contributions

A.M.W. and S.H.-B. conceived of the study. A.M.W. designed experiments with critical input from all authors. A.M.W. generated new reagents and performed experiments. A.M.W. and S.D. wrote data analysis software and analyzed data. A.M.W., S.D., E.M., and S.H.-B. interpreted data. A.M.W. prepared figures. A.M.W. and S.H.-B. wrote the manuscript with editorial input from all authors.

## Acknowledgements

We thank members of the Horne-Badovinac and Munro labs, Allison Zajac, Sherzod Tokamov, Ellie Heckscher, Michael Glotzer, and Carmen Williams for feedback throughout the study and comments on the manuscript. This work was supported by NIHR01GM126047 to S.H.B., NIHRO1HD88831 to E.M., NIHT32 HD055164 to A.M.W., and postdoctoral fellowships from the Chicago Fellows Program and Jane Coffin Childs Memorial Fund for Medical Research to S.D.

## Methods and Materials

### Materials, data, and code availability

New plasmids and *Drosophila* lines reported in this paper are available upon request to the corresponding author. The code necessary to reproduce core aspects of data analysis, along with example datasets, are available at *https://github.com/a9w/Fat2_polarizes_WAVE*. Additional data and code are available upon request to the corresponding author.

### *Drosophila* sources, care, and genetics

The sources and references of all stocks used in this study are listed in Supplemental Table 1 and the genotypes of *Drosophila* used in each experiment and associated figure panels are listed in Supplemental Table 2. *Drosophila* were raised at 25°C and fed cornmeal molasses agar food. Females 0-3 days post-eclosion were aged on yeast with males prior to dissection. In most cases, they were aged for 2-3 days at 25°C. Temperatures and yeasting times used for each experiment are reported in Supplemental Table 3. In all RNAi experiments, *traffic jam*>Gal4 (*tj*>Gal4)^53^ was used to drive RNAi expression in follicle cells and not in germ cells. Sra1-GFP and*fat2* mosaic epithelia were generated using the Flp/FRT method^54,55^, using FRT82B and FRT80B recombination sites, respectively. In both cases *tj*>Gal4 was used to drive expression of UAS>Flp recombinase.

### Generation of Sra1-GFP

Endogenous Sra1 was tagged C-terminally with enhanced GFP (GFP) following the general approaches described by Gratz *et al*. (2013) and Gratz *et al*. (2014)^56,57^. The guide RNA target sequence 5’-<u>GCTTAAATGCATCC CTTTCC</u>**GGG**-3’ was chosen with flyCRISPR Target Finder^57^. The underlined sequence was cloned into the pU6-BbsI-chiRNA plasmid, and the bold sequence is the adjacent PAM motif. For homologous recombination, homology arms approximately 2 kb long flanking the insertion target site were amplified from genomic DNA from the y1 M〈nos-Cas9.P〉ZH-2A w* (nanos-Cas9)^58^ background. GFP was amplified from the pTWG plasmid. A linker with sequence encoding the amino acids ‘GSGGSGGS’ was added to the N-terminal side of GFP. Homology arms, linker, and GFP were inserted into donor plasmid pDsRed-attP, which contains 3xP3-DsRed flanked by loxP sites for insertion screening and subsequent removal. The linker-GFP insertion was made immediately before the Sra1 stop codon. Guide and homologous recombination plasmids were injected by Genetivision Inc. into the nanos-Cas9 background. F1 males were screened for 3xP3-DsRed and then 3xP3-DsRed was excised by crossing to Cre-expressing flies (MKRS hsFLP/TM6b Cre).

### Egg chamber dissection

Ovaries were dissected into live imaging media (Schneider’s *Drosophila* medium with 15% fetal bovine serum and 200 *μ*g/mL insulin) in a spot plate using 1 set of Dumont #55 forceps and 1 set of Dumont #5 forceps. Ovarioles were removed from the ovary and from ovariole muscle sheathes with forceps. For live imaging, egg chambers older than the egg chamber to be imaged were removed from the ovariole strands by cutting through the stalk with a 27-gauge hypodermic needle. For fixed imaging, egg chambers older than stage 9 were removed prior to fixation. Removal of older egg chambers allows more compression of the imaged egg chamber between the slide and coverslip so that the basal surface of a field of cells can be imaged in a single plane. For a more detailed description and movies of dissection, see Cetera *et al*. (2016)^59^.

### Live imaging sample preparation

Following dissection, ovarioles were transferred to a fresh well of live imaging media. For membrane staining, CellMask Orange or Deep Red plasma membrane stain (Thermo Fisher Scientific, Waltham, MA, 1:500) was added and ovarioles incubated for 15 minutes, followed by a wash in live imaging media to remove excess stain before mounting. Ovarioles were then transferred to a glass slide with 20 μL of live imaging media. For CK-666 treatment, following plasma membrane staining, ovarioles were transferred to live imaging media with 750 μM CK-666 (Millipore Sigma, St. Louis, MO) and then mounted in the same media. Glass beads with diameter 51 μm were added to support the 22×22 mm #1.5 coverslip and limit egg chamber compression. Coverslip edges were sealed with melted petroleum jelly to prevent evaporation while imaging. Samples were checked for damage using the membrane stain or other fluorescent markers as indicators, and excluded if damage was observed. Slides were used for no more than 1 hour.

### Immunostaining and F-actin staining

Following dissection, ovarioles were fixed in 4% EM-grade formaldehyde in PBT (phosphate buffered saline + 0.1% Triton X-100) and then washed 3×5 minutes in PBT at room temperature. Egg chambers were incubated with primary antibodies in PBT overnight at 4°C (anti-Scar, 1:200) or for 2 hours at room temperature (anti-Discs Large, 1:20) while rocking. Ovarioles were then washed 3×5 minutes in PBT and incubated in secondary antibody diluted 1:200 in PBT for two hours at room temperature while rocking. F-actin staining was performed using either TRITC phalloidin (Millipore Sigma, 1:250) or Alexa Fluor 647 phalloidin (Thermo Fisher Scientific, 1:50). If TRITC phalloidin was the only stain or antibody used, it was added directly to the fixation media for 15 minutes of staining concurrent with fixation. Otherwise, TRITC phalloidin was added for 15-30 minutes at room temperature as the final staining step. Alexa Fluor 647 phalloidin staining was performed for two hours at room temperature while the sample was rocking, concurrent with secondary antibody staining where applicable. Ovarioles were then washed 3×5 minutes in PBT and mounted in 40 *μ*L SlowFade Diamond antifade on a slide using a 22×50 mm #1.5 coverslip, sealed with nail polish, and stored at 4°C until imaged.

### Microscopy

#### Laser scanning confocal microscopy

Laser scanning confocal microscopy was used for all fixed imaging and for live imaging of membrane-dyed egg chambers. Imaging was performed with a Zeiss LSM 800 upright laser scanning confocal with a 40x/1.3 NA EC Plan-NEOFLUAR oil immersion objective or a 63x/1.4 NA Plan-APOCHROMAT oil immersion objective, diode lasers (405, 488, 561, and 640 nm), and GaAsP detectors. The system was controlled with Zen 2.3 Blue acquisition software (Zeiss). Imaging was performed at room temperature. All images show the basal surface of stage 6-7 egg chambers except for Fig. S5A, bottom row, which shows follicle cells in cross-section. Cross-section images were used for egg chamber staging throughout. Laser scanning confocal microscopy was used to acquire the data in Figs. 2; 3B-F; 4A,E; 5A,C-G; S1; S2; S3A-C,E,F; S4; S5; S6; S7A; S8; Movies 1; 3; 4; 5; 6; 10.

#### TIRF microscopy

Near-TIRF microscopy was used to visualize Fat2-GFP, Sra1-GFP, Abi-mCherry, and F-Tractin-tdTomato^60^ dynamics at the basal surface. Near-TIRF imaging was performed with a Nikon ECLIPSE-Ti inverted microscope with Ti-ND6-PFS Perfect Focus Unit, solid-state 50 mW 481 and 561 nm Sapphire lasers (Coherent technology), motorized TIRF illuminator, laser merge module (Spectral Applied Research), Nikon CFI 100x Apo 1.45 NA oil immersion TIRF objective with 1.5x intermediate magnification, and Andor iXon3 897 electron-multiplying charged-coupled device (EM-CCD) camera. Image acquisition was controlled using MetaMorph software. For two color imaging, frames were collected for each color consecutively with the TIRF illumination angle adjusted in between. Imaging was performed at room temperature. For display, movies were corrected for bleaching using the histogram matching method in Fiji (ImageJ)^61,62^. TIRF microscopy was used to capture the data in Figs. 4B,D; 5B; S3D; S7B,C; S9; Movies 2; 7; 8; 9; 11; 12.

### Cell and protrusion segmentation from timelapses of cell membrane

Protrusions from timelapse datasets of the follicle cell basal surface stained with CellMask Orange (see Live imaging sample preparation) were segmented with the Python scikit-image and scipy libraries (Fig. S1)^63,64^. First, each cell was segmented and tracked, with manual corrections to cell-cell interface placements made using napari^65^. Next, a watershed-based approach was used to segment the regions of high fluorescence intensity at the interface of each pair of neighboring cells. This segmented shape encompasses the cell-cell interface and any associated protrusions from either neighboring cell. Last, to assign protrusions to the cell from which they originated, the segmented region was divided in two by the shortest path between its bounding vertices that lay entirely within the region. This approximates the position of the interface between the cells, and in subsequent steps we will call this line “the interface”. Each of the two resulting protrusion shapes was assigned as originating from the cell on the opposite side of the interface, because protrusions extend from one cell and overlap the other. Using this approach, all of the protrusive structures that emerge from one cell, and that overlap a single neighboring cell, are grouped together as a single segmented region for subsequent analysis.

### Measurement of membrane protrusivity, protrusion length, and protrusion orientation

After cell edges and associated protrusions were segmented, they were categorized as either protrusive or non-protrusive and their lengths and orientations using Python scikit-fmm, scikit-image, and scipy libraries. We use the term “membrane extensions” to refer to the cell edge shapes before the protrusive ones have been identified.

To measure the length of a membrane extension, we used two different metrics, each of which provides a single length value per cell edge. In one, we calculated the “average length” of a membrane extension as the membrane extension’s area divided by the length of the interface it extended across. As an alternate length measurement, we calculated its “longest length.” To do so, we first found its “tip”, defined as the pixel within the segmented region farthest from any point along the interface. We then found its “base”, the pixel along the interface that was closest to the tip. We defined membrane extension longest length as the length of the shortest path between base and tip that lay entirely within the membrane extension. To categorize membrane extensions as *protrusive* or *non-protrusive* throughout the study, we used the “average length” metric. We measured the average length distribution in CK-666-treated epithelia, which are nearly non-protrusive and so provided a measure of the width of the cell-cell interface alone. For all conditions, we categorized a membrane extension as protrusive if its average length was greater than the 98th percentile of length of CK-666-treated epithelia. We then defined the protrusivity of an entire epithelium as the ratio of protrusive to total cell edges in the field of view. We also report two alternate measurements of the protrusivity of an epithelium. In one, We calculate epithelial protrusivity as above, but substitute the longest length as our length measurement (Fig. S2A). In a second, cutoff-independent epithelial protrusivity measurement, we report the epithelium-mean average membrane extension length (Fig. S2B). Swarm plots of each of these analyses were generated using GraphPad Prism 9 (GraphPad, San Diego, CA), as were all other swarm plots.

For analysis of protrusion orientation, we included only the membrane extensions categorized as protrusive according to the “average length” metric. We defined a protrusion’s orientation as the orientation of the vector from its base to its tip. Polar histograms, generated in Python with matplotlib^66^, show the distribution of protrusion orientations. In these plots, bar area is proportional to the number of protrusions in the corresponding bin.

### Quantification of F-actin and Sra1-GFP cell-cell interface and non-interface basal surface fluorescence

Cells and cell-cell interfaces were segmented as described above. Cells and interfaces in contact with the tissue border or image border were excluded from analysis. For interface fluorescence intensity, interfaces were dilated by 5 pixels, and mean fluorescence intensity calculated from within this region. Non-interface basal surface fluorescence intensity was calculated as the mean of the remaining (non-interface) tissue surface. For F-actin cell-cell interface enrichment measurements, the overall brightness of the phalloidin staining varied between epithelia independent of genotype. To control for this variation we subtracted the mean intensity of the epithelium’s non-interface basal surface from its mean interface intensity measurement. This value, the degree of F-actin interface enrichment, was used as a proxy for F-actin protrusivity.

### Quantification of F-actin and Sra1-GFP planar polarity

As a simple planar polarity measurement, we quantified mean F-actin (phalloidin) or Sra1-GFP fluorescence intensity along each cell-cell interface as a function of the interface’s orientation. To do this, cells and cell-cell interfaces were segmented as described above. For interface angle measurements, the angular distance between the line defined by the interface-bounding vertices and the anterior-posterior (horizontal) axis was calculated. For interface fluorescence intensity measurements, interface regions were identified as segmented interfaces dilated by 5 pixels. Vertices, dilated by 10 pixels, were excluded from interface regions. Mean fluorescence intensity was calculated within each interface region, and background (the mean non-interface basal surface fluorescence intensity of all cells in the image) was subtracted. To calculate the leading-trailing interface enrichment of F-actin or Sra1-GFP for each egg chamber, interface fluorescence intensities were averaged for all interfaces with angles between 0° and 10° (leading-trailing interfaces), and between 80° and 90° (side interfaces). The leading-trailing interface enrichment is the ratio of these numbers.

### Autonomy analysis in mosaic epithelia

Egg chambers were stained with Alexa Fluor 647 phalloidin to mark protrusions, which indicate migration direction, and to determine whether egg chambers were planar polarized. We analyzed only S6-7 egg chambers with mixtures of control and *fat2* cells that had global stress fiber alignment orthogonal to the anterior-posterior axis, indicating global planar polarity. Since migration is required to maintain planar polarity^16^, this also indicates that the epithelium was migratory. Looking at phalloidin and genotype markers only, we drew 10 pixel-wide segmented lines along leading-trailing interface boundaries of different genotype combinations in Fiji. Lines were drawn along all visible, in-focus *fat2-control* and *control-fat2* boundaries and a similar number of control-control and *fat2-fat2* boundaries. A boundary category was excluded if there were fewer than 3 usable interfaces to measure. Mean Sra1-GFP fluorescence intensities were calculated for each interface type in each egg chamber. For a diagram of this method, see Barlan *et al*. (2017)^15^. To quantify non-interface basal surface fluorescence, we drew polygonal regions of the basal surface of control and *fat2* cells, excluding cells immediately behind those of a different genotype. Egg chambers were excluded if there were fewer than 3 usable cells of either genotype. Mean Sra1-GFP fluorescence intensities were calculated within these polygonal regions for all control cells and all *fat2* cells in an egg chamber.

### Quantification of migration rate

Egg chambers were dissected, dyed with CellMask Orange, and mounted for live imaging as described above. Several ovarioles were mounted on each slide, with each ovariole terminating in a S6-7 egg chamber. Timelapse imaging was performed for 30 minutes with frames acquired every 30 seconds. Multi-point acquisition was used to obtain movies of up to 5 egg chambers simultaneously. To generate a kymograph, a line was drawn along the axis of migration at the center of the anterior-posterior egg chamber axis in Fiji. In these kymographs, cell-cell interfaces are visible as lines, and their slope gives a measurement of cell migration rate. Egg chamber migration rates were calculated from the average of 4 cell interface slopes from each kymograph. Egg chambers that clearly slowed down over the course of the timelapse, visible as curvature in the interface lines in the kymographs, were excluded. For an illustration of this method, see Barlan *et al*)^15^.

### Cell perimeter kymograph generation and interpretation

To visualize the distribution of Sra1-GFP along cell-cell interfaces over time, we generated kymographs of cell perimeters from timelapses of Sra1-GFP-expressing epithelia obtained using near-TIRF microscopy. Perimeters were drawn manually in Fiji in each frame with the pencil tool, and then these perimeters were used to generate kymographs in Python. Perimeters were thinned to 1 pixel and then perimeter pixels were sequenced with Python scikit-image and scipy libraries. Kymographs were generated with matplotlib. Kymograph rows were constructed by linearizing the perimeters from each frame, starting with the pixel directly above the cell centroid (the center of the trailing edge in control cells) and continuing counter-clockwise. Each row shows the fluorescence intensity of the perimeter pixels in sequence. Cell perimeter lengths varied between frames, so kymograph row lengths varied and were aligned to their center position.

At the spatial and temporal resolution of the timelapses and corresponding kymographs, we cannot evaluate differences in the dynamics the puncta-scale WAVE complex accumulations highlighted in Fig. 5. Instead, we focus on the “region”-scale distribution of Sra1-GFP, and the stability of that distribution over time. The regions we refer to here are approximately the length of a cell-cell interface, with variation. Because the kymographs are generated from epithelia in which all cells express Sra1-GFP, we need additional information to identify the cell to which a region of Sra1-GFP enrichment belongs. We infer that Sra1-GFP is predominantly at leading edges in polarized, migratory epithelia based on the Sra1-GFP distribution in epithelia with mosaic Sra1-GFP expression (Fig. 3B). Based on consistent correlation between Sra1-GFP enrichment and the presence of protrusions (Fig. 3B, 4A, S7A), and its known role building lamellipodia as part of the WAVE complex^17,18,41^, we also infer that regions of Sra1-GFP enrichment belong to the cell that is protruding outward regardless of genotype. Our interpretations of Sra1-GFP enrichment patterns in movies and corresponding kymographs are made with these assumptions.

### Colocalization of proteins along the leading-trailing interface

Data used for colocalization analysis were collected with 63x/1.4 NA Plan-APOCHROMAT oil immersion objective to minimize chromatic aberration. Linescans were generated in Fiji by manually drawing a 10 pixel-wide segmented line along rows of leading-trailing interfaces at the follicle cell basal surface. At least 20 leading-trailing interfaces were included per egg chamber. For the Fat2^ΔICD^ condition, in which the distribution of Fat2 expands beyond leading-trailing interfaces, we measured colocalization either along randomly oriented interfaces (Fig. 5E) or leading-trailing interfaces (Fig. S8B) and obtained very similar results. Fluorescence intensities along the linescans were obtained with the PlotProfile function, which averages pixel intensities along the width of the line and reports a list of averaged values along the line’s length. Spearman’s correlation coefficients were calculated for each egg chamber in Python with the scipy.stats module. Failure to exactly follow leading-trailing interfaces and cusps in the segmented lines will artificially inflate the measured correlation, so we used correlation between E-cadherin-GFP^67^ and Abi-mCherry as a negative control that is also subject to this inflation. Abi-mCherry and E-cadherin-GFP are slightly displaced from each other (anticorrelated) along the length of protrusions (the width of the linescans), but averaging across the line width collapses this displacement, resulting in measured intensity signals that are roughly uncorrelated. Spearman’s correlation coefficients ± standard deviation are reported in the text. Linescans of leading-trailing interfaces were plotted using the fluorescence intensities from along the leading-trailing interfaces of two cells. Intensities from each fluorophore were rescaled between 0 and 1 and plotted with matplotlib in Python.

### Protrusion profile generation

Viewing only the F-actin channel in Fiji, we drew 1 pixel-wide lines down the length of F-actin bundles at the leading edge. Fluorescence intensities along these lines were obtained for all fluorophores with the Fiji PlotProfile function. In Python, these traces were aligned to the pixel with highest Fat2-3xGFP or Ena-GFP intensity (Fig. 5G, S8F). To calculate standard deviation, all traces were first rescaled individually so that their values ranged between 0 and 1. To plot “protrusion profiles,” the mean fluorescence was determined for each fluorophore at each pixel position, and then average values were rescaled between 0 and 1. Plots of protrusion profiles were generated with matplotlib.

### Movie generation

Migration motion was subtracted from several timelapse movies of migratory cells or epithelia for ease of visualization. Motion subtraction was performed using the Fiji MultiStackReg plugin “translation” transformation^68^(control condition in Movies 1; 6; 8) or by aligning to the centroid of a tracked cell in each frame using the scikit-image library (Movies 7; 10). Labels were added to movies in Fiji and then exported as uncompressed .avi files. These were encoded as 1080p30 .mp4 files with H.264 (x264) video encoder using HandBrake 1.4.

### Reproducibility and statistical analysis

Visibly damaged egg chambers were excluded from all analyses. Each experiment was performed at least two independent times, and results confirmed to be qualitatively consistent. Each experiment included egg chambers pooled from multiple flies. Experiments and analysis were not randomized or performed blinded. Sample sizes were not predetermined using a statistical method. The number of biological replicates (n), statistical tests performed, and their significance can be found in figures or figure legends. Based on visual inspection, all data on which statistical tests were performed followed an approximately normal distribution, so tests assuming normalcy were used. Alpha was set to 0.05 for all statistical tests. Paired statistical tests were used for comparisons of cells of different genetic conditions within mosaic epithelia, except if all epithelia did not have all genetic conditions represented, in which case an unpaired test was used so that all samples could still be included. All t-tests were two-tailed. One-sample t-tests were used when comparing a distribution of ratios to a null expectation of one. A one-way ANOVA was used when multiple pairs of conditions were compared, with the exception of plots in Figs. 2C, S2A, and S3C, for which the variance did not appear consistent between conditions, so Welch’s ANOVA was used instead. For post-hoc comparison tests, all pairs of conditions present in the corresponding plot were compared using post-hoc Tukey’s multiple comparisons test with the following exception: the data plotted in Figs. 5E and S8B were analyzed together, and all conditions were compared to Fat2-Abi and E-cadherin-Abi only, and in Fig. S6C only data from the same region (total, interface, or non-interface) was compared. For these, Šidák’s multiple comparisons tests were used. For Welch’s ANOVA, Dunnet’s T3 multiple comparisons tests were used. P-values reported for all post-hoc tests were adjusted for multiple comparisons. All statistical tests except for the calculation of Spearman’s correlation coefficients were performed in GraphPad Prism 9.

---

**Figure S1:**
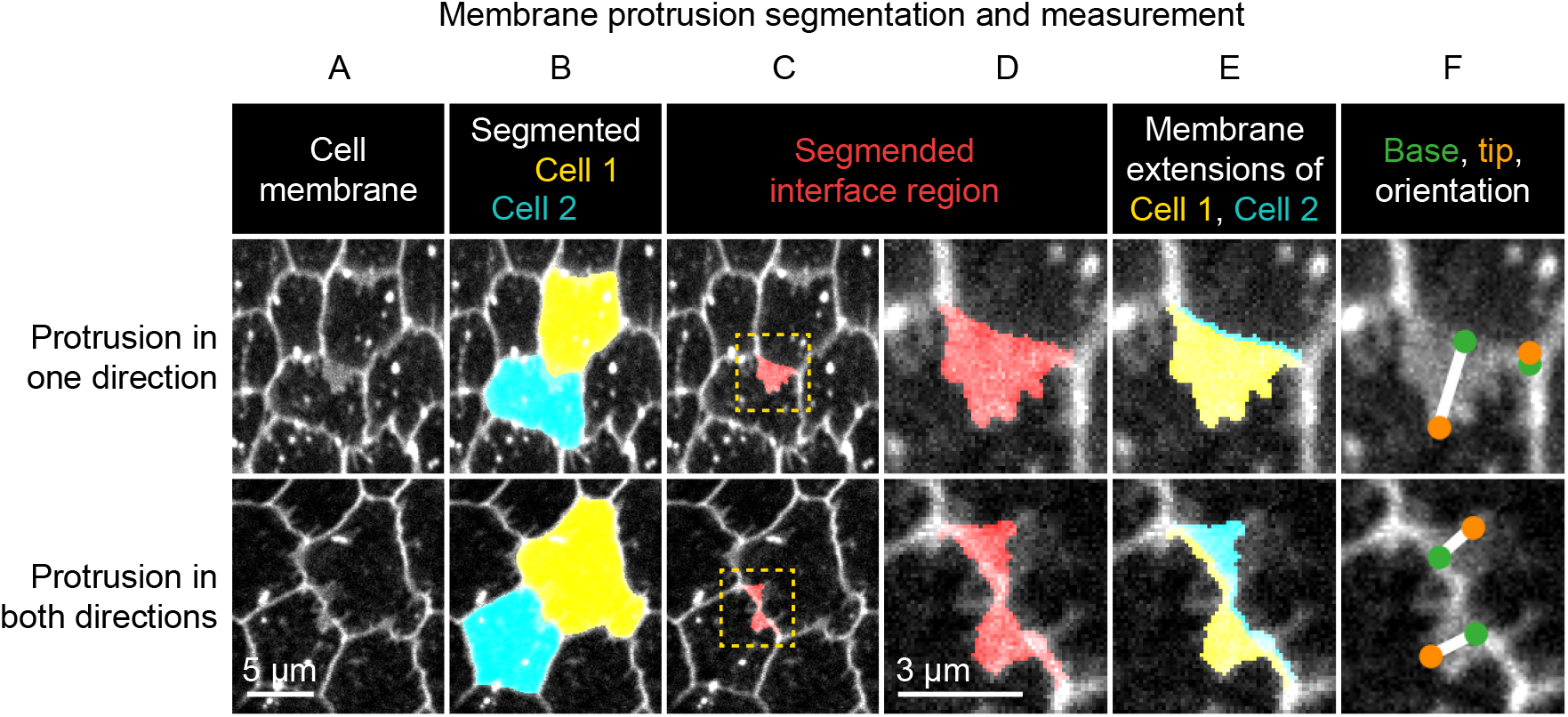
Method used to segment and measure membrane protrusions. Top row shows an example of a pair of neighboring cells in which one cell is protruding across their shared interface. Bottom row shows a case in which both cells are protruding across the interface. **A**, Cell interfaces and protrusions were labeled with a membrane dye and timelapses of the basal surface were collected. **B**, Cells were automatically segmented with a watershed-based method, and segmentation errors were hand-corrected. **C**, The bright interface region between each pair of neighboring cells was identified using a watershed-based method. This region includes the interface and any membrane protrusions that extend across it. **D**, An enlargement of the boxed regions of (C). **E**, The interface region was divided into two parts by the shortest path from vertex to vertex within the region, which approximates the true cell-cell interface position. The two resulting regions were then assigned to the cell from which they each extended. The area of these regions and the length of the interface between them was used to define average membrane extension length (as described in Methods). **F**, The tip and base of each region were identified, and then used to measure lengths and orientations (see Methods). Associated with Fig. 2.

**Figure S2:**
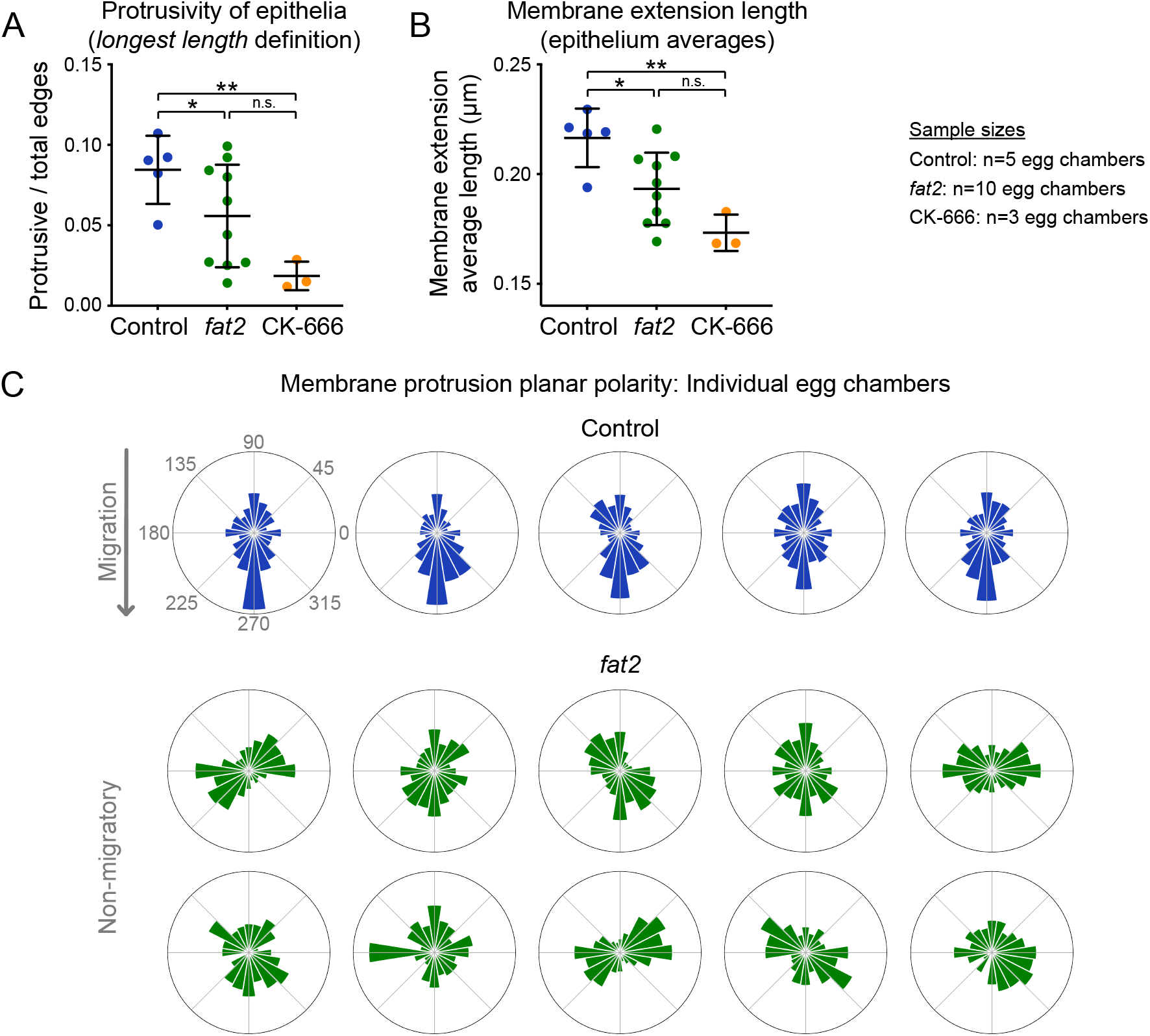
Membrane extension length and protrusion orientation in individual egg chambers. **A**, Plot showing the ratio of protrusive to total edges, with protrusivity defined in terms of membrane extension *longest length* (see Methods). In agreement with the *average length* definition, the protrusivity of *fat2* epithelia is variable, with a distribution overlapping with control and CK-666-treated epithelia. Welch’s ANOVA (W(2,8.74)=19.3, p=0.0006) with Dunnet’s T3 multiple comparisons test; n.s. p=0.16, *p=0.02, **p=0.0024. **B**, Plot showing the mean membrane extension lengths of control, textitfat2, and CK-666-treated egg chambers, with membrane extension length defined as *average length* (see Methods). With this cutoff-independent protrusivity measurement, the protrusivity of *fat2* egg chambers is intermediate between control and CK-666, with a wider distribution that overlaps both. Welch’s ANOVA (W(2,7.31)=14.75, p=0.0027) with Dunnet’s T3 multiple comparisons test; n.s. p=0.069, *p=0.042, **p=0.0036. **C**, Polar histograms showing the distribution of membrane protrusion orientations in individual control and *fat2* egg chambers. Anterior is left, posterior is right, and images were flipped as needed so that migration is downward for control epithelia, in which membrane protrusions are biased in the direction of migration. In *fat2* epithelia, protrusions have varying levels of axial bias and little or no vectorial bias. Associated with Fig. 2.

**Figure S3:**
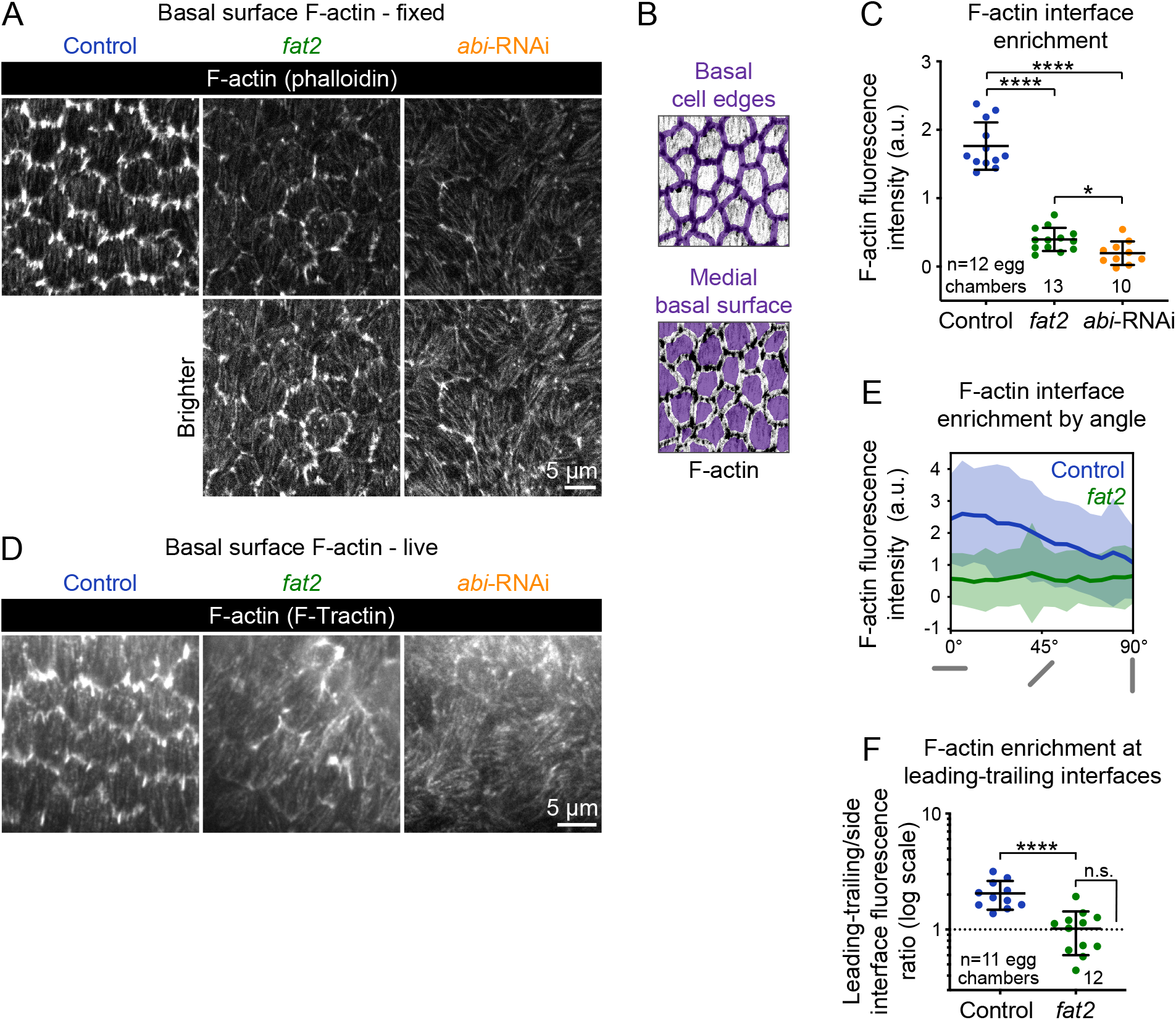
Actin protrusions are reduced and unpolarized without Fat2 and further reduced without the WAVE complex. **A**, Images showing phalloidin staining of F-actin in control, *fat2*, and *abi*-RNAi-expressing epithelia. Bottom row shows the same images with displayed brightness increased. From dataset quantified in C,E,F. **B**, Examples of segmented cell-cell interfaces or medial basal surfaces overlayed on F-actin. **C**, Plot of the difference in F-actin fluorescence intensity between cell interfaces and medial basal surfaces shows that while *fat2* and *abi*-RNAi epithelia have less F-actin interface enrichment than control epithelia, F-actin interface enrichment remains higher in *fat2* epithelia than *abi*-RNAi epithelia. Welch’s ANOVA (W(2, 19.84)=94.68, p<0.0001) with Dunnet’s T3 multiple comparisons test; *p=0.033, ****p<0.0001. **D**, Frames from timelapse movies of *control,fat2*, and *abi*-RNAi epithelia with F-actin labeled with F-Tractin-tdTomato. As with phalloidin staining, the protrusivity of *fat2* epithelia is intermediate between that of control and *abi*-RNAi epithelia. Brightness display settings vary between genotypes to correct for variability in F-Tractin-tdTomato expression levels. See related Movie 2.**E**, Plot of F-actin fluorescence intensity at cell interfaces as a function of interface angular distance from horizontal. Gray bars below the x-axis represent interface angles. **F**, Plot of the F-actin fluorescence ratio between leading-trailing (0-10°) and side (80-90°) interfaces, a measure of F-actin enrichment along leading-trailing interfaces. F-actin is enriched at leading-trailing interfaces in control, but not *fat2*, egg chambers. Control-*fat2* comparisons: unpaired t-test, ****p<0.0001. Comparison between *fat2* and one (dashed line, the expectation if there is no enrichment): one sample t-test, n.s. p=0.88. **C,E,F**, Bars (C,F) or lines and shaded regions (E) indicate mean ±SD. Associated with Fig. 2; Movie 2.

**Figure S4:**
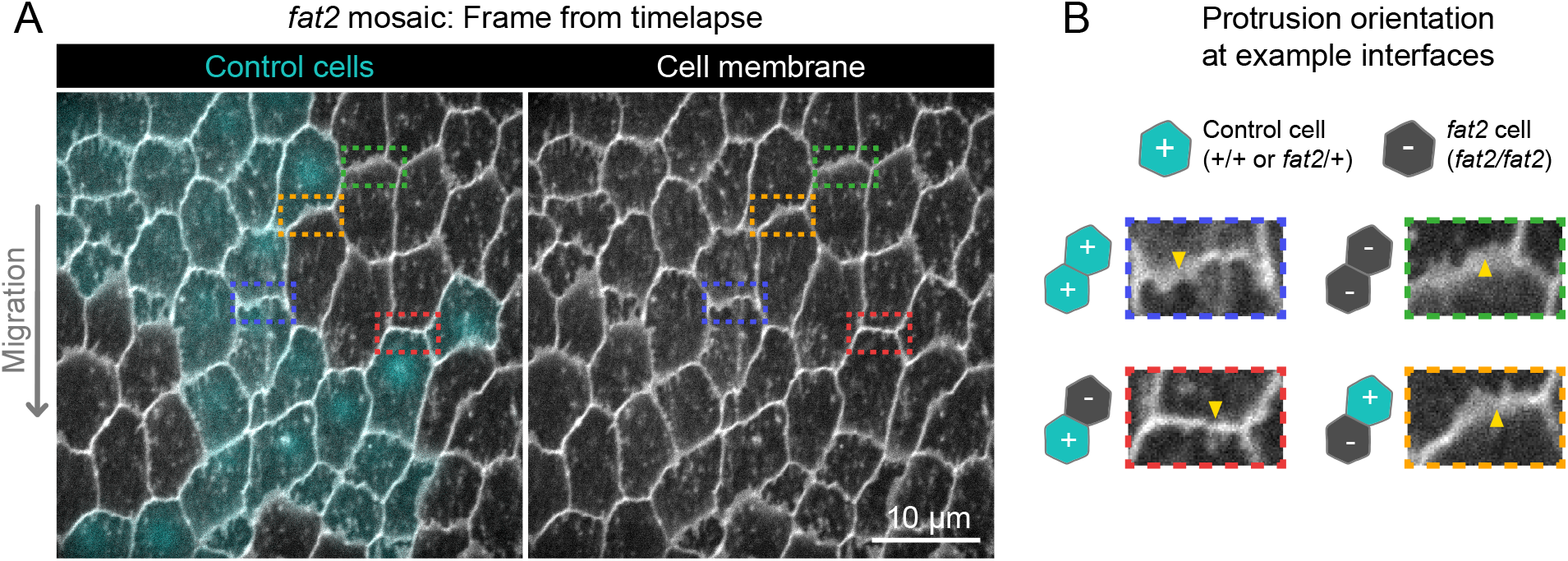
Fat2 acts locally across the cell interface to orient membrane protrusions. **A**, Timelapse frame of a *fat2* mosaic epithelium with cell membrane labeled, used to evaluate protrusion orientations in control or *fat2* cells within a migratory context. Boxes indicate examples of leading-trailing interfaces between neighbor pairs with each possible combination of genotypes. See related Movie 4.**B**, Larger images of the interfaces boxed in (A), showing that protrusions are misoriented when *fat2* cells are ahead of the interface regardless of the genotype of the cell behind the interface. Arrows point in the direction of protrusion. Associated with Fig. 2; Movie 4.

**Figure S5:**
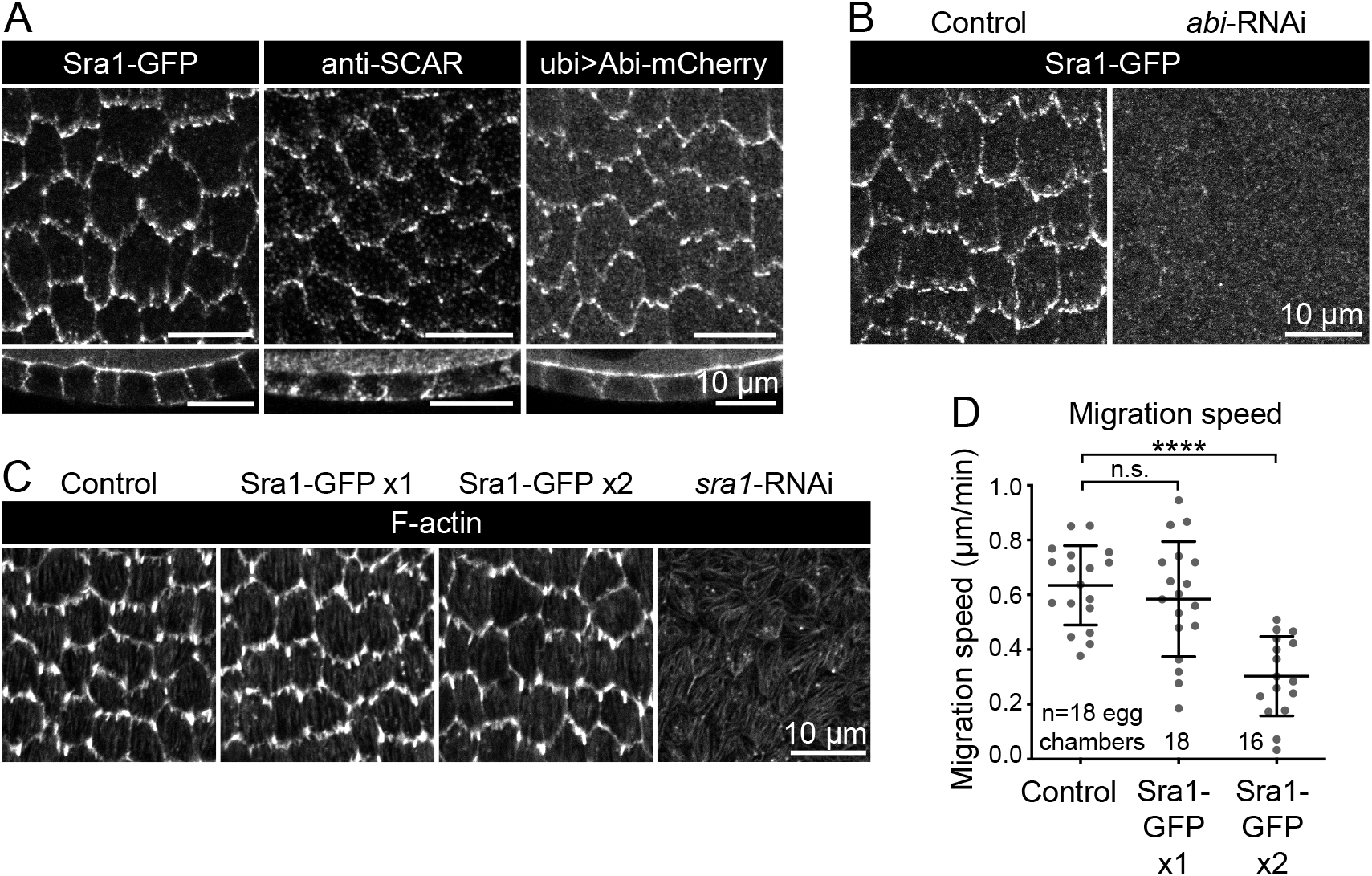
Evaluation of endogenous Sra1-GFP functionality. **A**, Images comparing the localization of markers ofWAVE complex subunits: Sra1-GFP, Scar antibody, and Abi-mCherry, at the basal surface (top row) and in cross-section (bottom row). **B**, Images of Sra1-GFP localization in control and *abi*-RNAi-expressing epithelia. Sra1-GFP is dispersed in the absence of Abi. **C**, Images showing phalloidin-stained F-actin in epithelia with wild-type Sra1, one or two copies of Sra1-GFP, or expressing *sra1*-RNAi, used to assess the appearance of protrusions in each condition. **D**, Plot of migration speed of epithelia with wild-type Sra1 or one or two copies of Sra1-GFP. Migration speed is reduced when both Sra1 copies are GFP-tagged. One-way ANOVA (F(2,49)=18.37, p<0.0001) with post-hoc Tukey’s test; n.s. 0.66, ****p<0.0001. See related Movie 5. Associated with Fig. 3; Movie 5.

**Figure S6:**
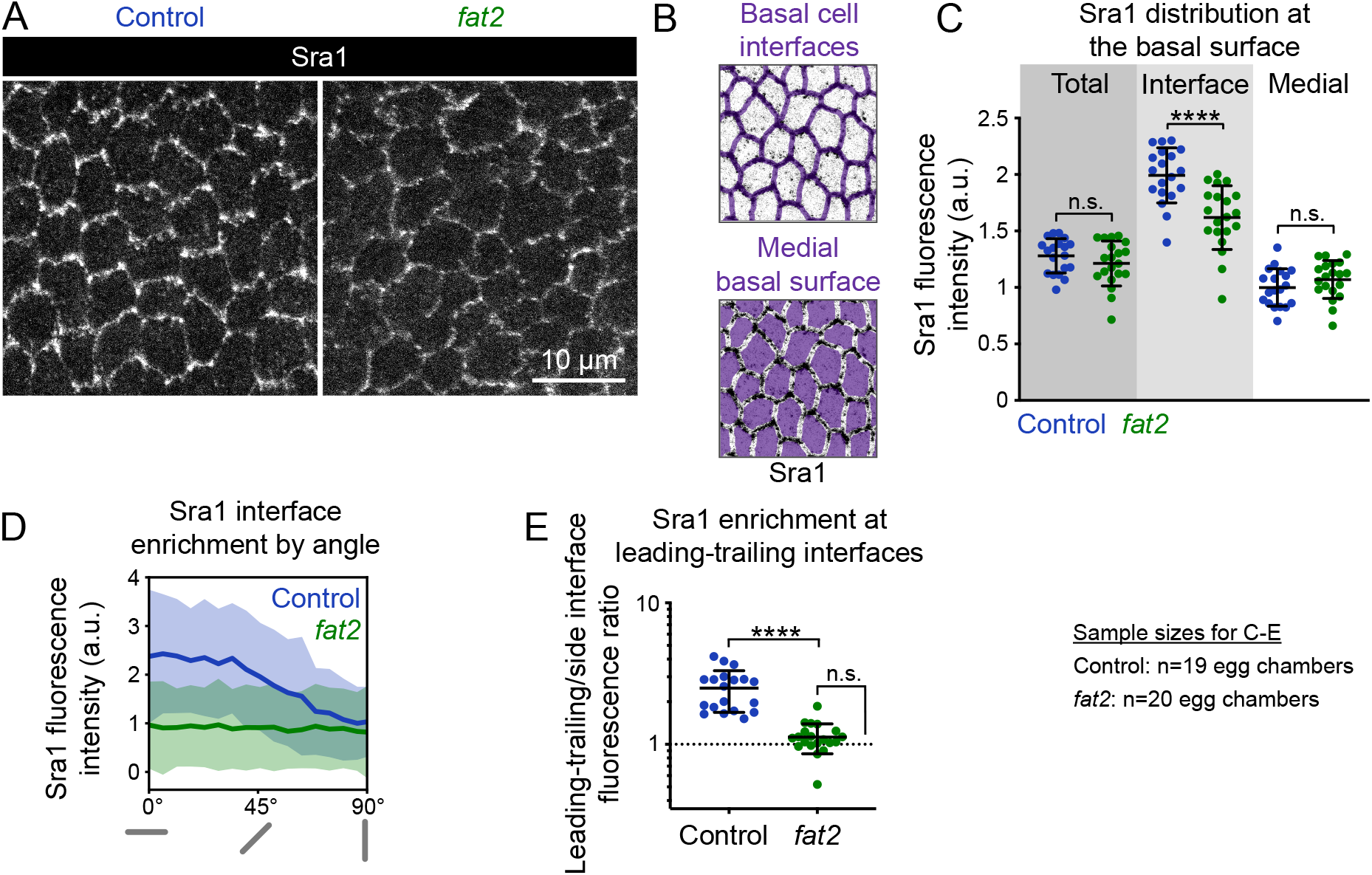
Fat2 concentrates the WAVE complex at cell-cell interfaces and polarizes it across the epithelium. **A**, Images of Sra1-GFP at the basal surface in control and *fat2* epithelia. **B**, Examples of segmented cell-cell interfaces or medial basal surfaces overlaid on Sra1-GFP images. **C**, Plot of mean Sra1-GFP fluorescence intensity across the entire basal surface (total), at cell-cell interfaces, and at medial basal surfaces in control and *fat2* epithelia. One-way ANOVA (F(5,111)=63.22, p<0.0001) with post-hoc Šidák’s test; n.s. (left to right) p=0.67, 0.64, ****p<0.0001. **D**, Plot of Sra1-GFP fluorescence at cell-cell interfaces as a function of interface angular distance from horizontal. Gray bars below the x-axis represent interface angles. **E**, Plot of the Sra1-GFP fluorescence intensity ratio between leading-trailing (0-10°) and side (80-90°) interfaces, a measure of Sra1-GFP enrichment along leading-trailing interfaces. Sra1-GFP is enriched at leading-trailing interfaces in control, but not *fat2*, epithelia. *Control-fat2* comparison: unpaired t-test, ****p<0.0001. Comparison between *fat2* and one (dashed line, the expectation if there is no enrichment): one sample t-test, n.s. p=0.052. Bars (C,E) or lines and shaded regions (D) indicate mean ±SD. Associated with Fig. 3.

**Figure S7:**
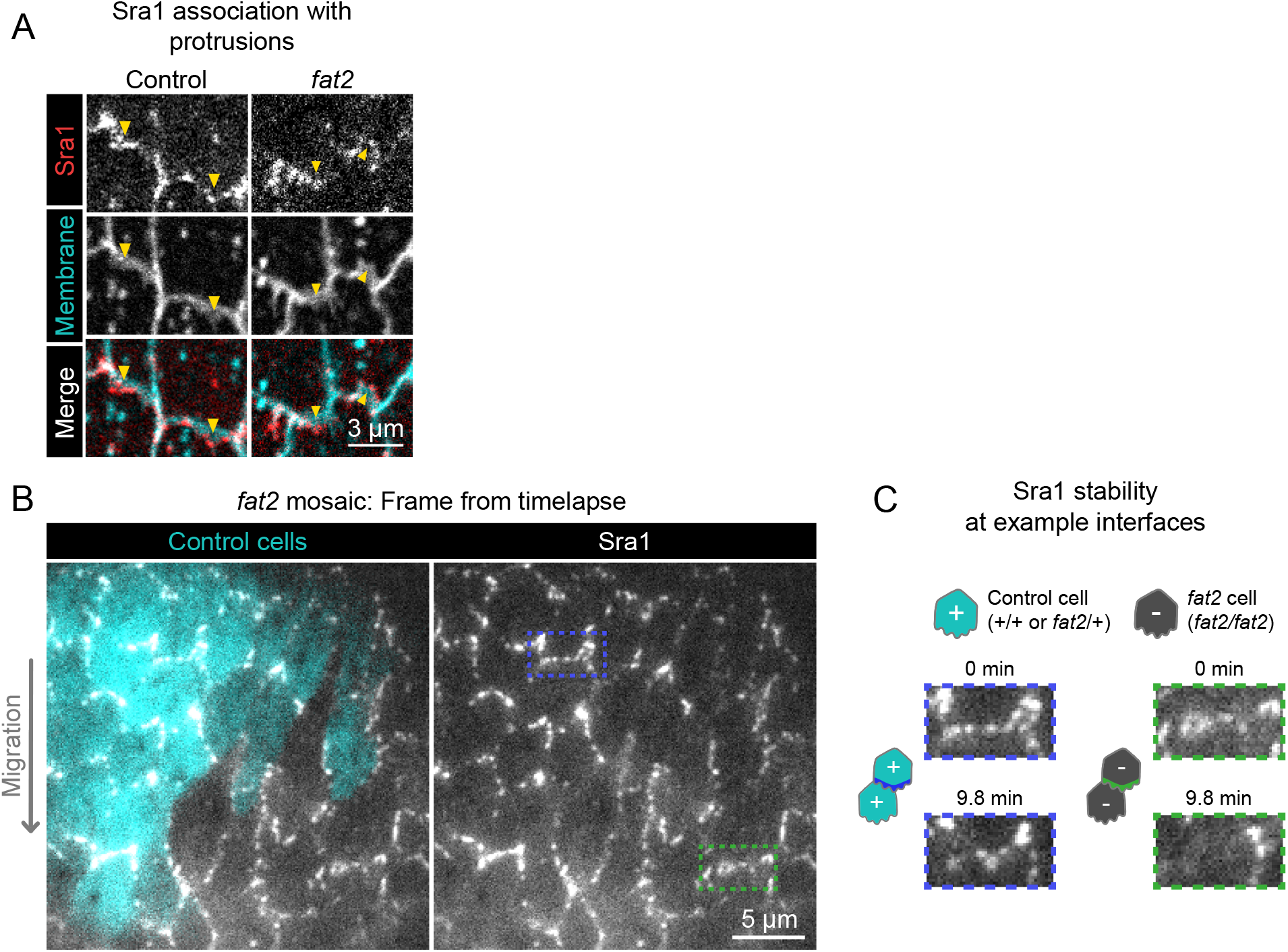
Fat2 stabilizes domains of WAVE complex enrichment locally across the cell-cell interface. **A**, Timelapse frames showing pairs of cell interfaces from control or *fat2* epithelia expressing Sra1-GFP and labeled with a membrane dye. Arrows indicate membrane protrusions. Sra1-GFP is enriched at protrusion tips in both control and *fat2* epithelia. See related Movie 6.**B**, Timelapse frame of a *fat2* mosaic epithelium in which all cells express Sra1-GFP, used to evaluate Sra1-GFP dynamics in control or *fat2* cells within a migratory context. Boxes indicate a leading-trailing interface between two control cells (blue) or *fat2* cells (green). See related Movie 9.**C**, Larger images of the interfaces boxed in (B), taken 9.8 minutes apart. Sra1-GFP is initially enriched along both interfaces. It remains enriched in the control interface throughout, but loses enrichment along the *fat2* interface. Associated with Fig. 4; Movies 6, 9.

**Figure S8:**
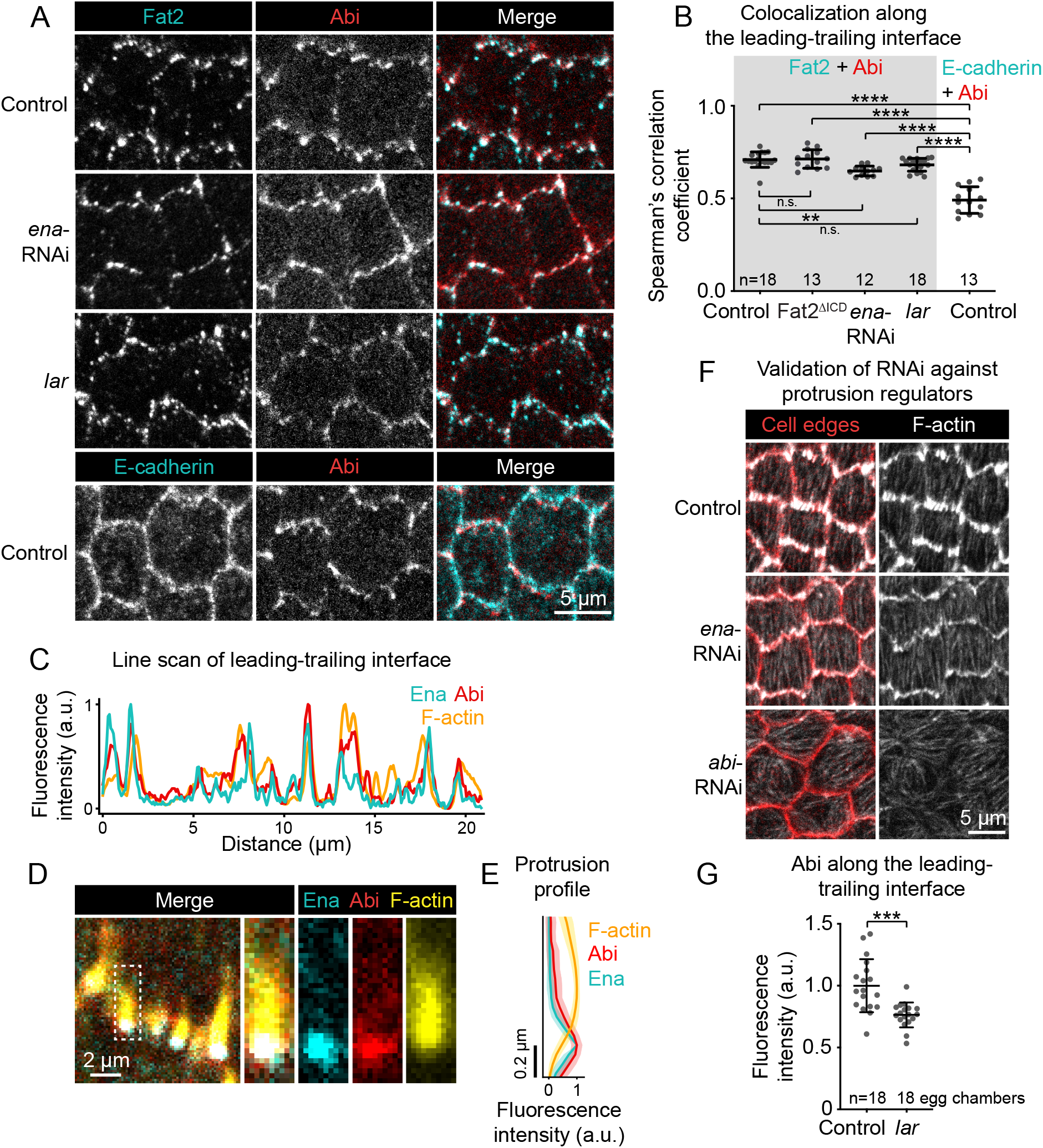
Ena and Lar are not required for colocalization between Fat2 and the WAVE complex. **A**, Images of cells expressing Fat2-3xGFP and Abi-mCherry in control, *lar*, and *ena*-RNAi backgrounds (top 3 rows) or E-cadherin-GFP and Abi-mCherry (bottom row, negative control for colocalization measurements). **B**, Plot of Spearman’s correlation coefficients of Abi-mCherry and Fat2-3xGFP (or Fat2^ΔIC^D-3xGFP) (gray background) or E-cadherin-GFP (white background) along leading-trailing interfaces show that Fat2 and Abi colocalize in all four conditions more strongly than E-cadherin and Abi. Bars indicate mean ±SD. One-way ANOVA (F(5,81)=44.86, p=0.0164) with post-hoc Tukey’s test; n.s. (left to right) p>0.99, p=0.41, **p<0.0046, ****p<0.0001. **A,B**, Control, Fat2-3xGFP, and Abi-mCherry images and Spearman’s coefficients are also in Fig. 5A,E. **C**, Line scan of GFP-Ena, Abi-mCherry, and F-actin (phalloidin) fluorescence intensity along a leading-trailing interface region, showing their corresponding peaks of enrichment. **D**, Image showing the GFP-Ena, Abi-mCherry, and F-actin (phalloidin) at the leading edge and in the boxed filopodium. **E**, Plot of mean fluorescence intensity of F-actin, Abi-mCherry, and GFP-Ena along the length of filopodia showing their relative distribution. Lines and shaded regions indicate mean ±SD. n=54 filopodia (used for SD), 39 cells from 2 egg chambers. **F**, Images of F-actin (phalloidin) and cell interfaces (anti-Discs Large) in control, *ena*-RNAi, and *abi*-RNAi backgrounds. Expression of *ena*-RNAi strongly depletes filopodia, and *abi*-RNAi expression removes both filopodia and lamellipodia. **G**, Plot of mean fluorescence intensity of Abi-mCherry along leading-trailing interfaces in control epithelia or similarly-oriented interfaces in *lar* epithelia, some of which are non-migratory. Associated with Fig. 5.

**Figure S9:**
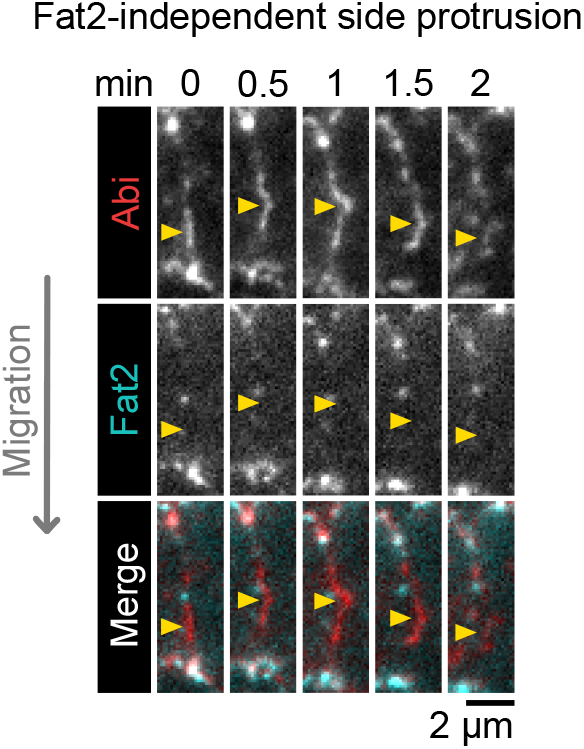
The WAVE complex occasionally accumulates at side-facing interfaces away from Fat2. Timelapse frames of a side-facing cell-cell interface from an epithelium expressing Abi-mCherry and Fat2-3xGFP. Arrows indicate a site of transient Abi-mCherry accumulation, protrusion, and dissipation with no corresponding Fat2-3xGFP enrichment. See related Movie 12. Associated with Fig. 5; Movie 12.

## Movie captions

Movie 1: **Membrane protrusivity of control, *fat2*, and CK-666-treated epithelia.** Control, *fat2*, and CK-666-treated epithelia labeled with a membrane dye. Bottom row shows segmented edges. Protrusive edges, defined as ones with with membrane extensions longer than the 98th percentile of those of CK-666-treated epithelia, are red. Non-protrusive edges are white. The control field of view moves to follow the cells as they migrate. Acquired with laser scanning confocal microscope. Associated with Fig. 2A.

Movie 2: **F-actin protrusivity and protrusion polarity of control and *fat2* epithelia.** Control, *fat2*, and *abi*-RNAi epithelia with F-actin labeled with F-Tractin-tdTomato. The protrusivity of *fat2* epithelia is intermediate between that of control and *abi*-RNAi epithelia. Brightness display settings vary between genotypes to correct for variability in F-Tractin-tdTomato expression levels. Acquired with TIRF microscope. Associated with Fig. S3D.

Movie 3: **Protrusion orientation in control and *fat2* epithelia.** Control and *fat2* epithelia are labeled with a membrane dye, and arrows indicating the orientation of protrusions are overlayed. Arrows originate at protrusion bases and have lengths proportional to protrusion lengths. Acquired with laser scanning confocal microscope. Associated with Fig. 2A.

Movie 4: **Membrane protrusion in a *fat2* mosaic epithelium.** A *fat2* mosaic epithelium with cell membrane labeled, used to evaluate protrusion orientations in control or *fat2* cells within a migratory context. Boxes indicate examples of leading-trailing interfaces between neighbor pairs with each possible combination of genotypes. Acquired with laser scanning confocal microscope. Associated with Fig. S4.

Movie 5: **Migration of epithelia with endogenously-tagged Sra1-GFP.** Epithelia with unlabeled Sra1 (Control), one copy of Sra1-GFP and one unlabeled Sra1, or two copies of Sra1-GFP, imaged at the mid-plane between apical and basal cell surfaces. Representative of timelapse movies used to measure migration speed. Acquired with laser scanning confocal microscope. Associated with Fig. S5D.

Movie 6: **Sra1 enrichment at protrusion tips in control and *fat2* epithelia.** Pairs of cell edges from control or *fat2* epithelia expressing Sra1-GFP and labeled with a membrane dye. Sra1-GFP is enriched at protrusion tips in both control and *fat2* epithelia. The control movie field of view moves to follow the cells as they migrate. Associated with Fig. S7A.

Movie 7: **WAVE complex-enriched domain dynamics in control and *fat2* cells.** Cells from control and *fat2* epithelia expressing Sra1-GFP. Laser intensity and brightness display settings differ between genotypes. Used to evaluate the stability of domains of Sra1-GFP accumulation. The control field of view moves to follow the cell as it migrates. Associated with Fig. 4B,D.

Movie 8: **WAVE complex-enriched domain dynamics in control *and fat2* epithelia.** Fields of cells from control and *fat2* epithelia expressing Sra1-GFP. The control movie field of view moves to follow the cells as they migrate. Laser intensity and brightness display settings differ between genotypes. Used to evaluate the stability of domains of Sra1-GFP accumulation. Wider view of the epithelia shown in Fig. 4B,D.

Movie 9: **WAVE complex-enriched domain dynamics in a *fat2* mosaic epithelium.** A *fat2* mosaic epithelium in which all cells express Sra1-GFP, used to evaluate WAVE complex-enriched domain dynamics in control or *fat2* cells within a migratory context. Associated with Fig. S7B,C.

Movie 10: **Dynamics of protrusive domains in control and *fat2* cells.** Top row shows the interfaces and membrane protrusions of one cell and its neighbors, labeled with a membrane dye. The segmented membrane extensions originating from the centered cell are overlaid in the bottom row. The control field of view moves to follow the cell as it migrates. Associated with Fig. 4E.

Movie 11: **Colocalization of puncta of Fat2 and the WAVE complex along leading-trailing interfaces.** The leading-trailing interfaces of two cells expressing Fat2-3xGFP and Abi-mCherry, used to compare the distributions of Fat2 and the WAVE complex over time. The Fat2-3xGFP channel is offset 2 pixels downward so puncta positions can be more easily compared. Associated with Fig. 5B.

Movie 12: **WAVE complex accumulation at side-facing protrusions away from Fat2.** A side-facing cell-cell interface from an epithelium expressing Abi-mCherry and Fat2-3xGFP. Arrows indicate a site of transient Abi-mCherry accumulation, protrusion, and dissipation with no corresponding Fat2-3xGFP enrichment, in contrast to the colocalized Fat2-3xGFP and Abi-mCherry on the leading-trailing interfaces above and below. Associated with Fig. S9.

**Supp. Table 1:**
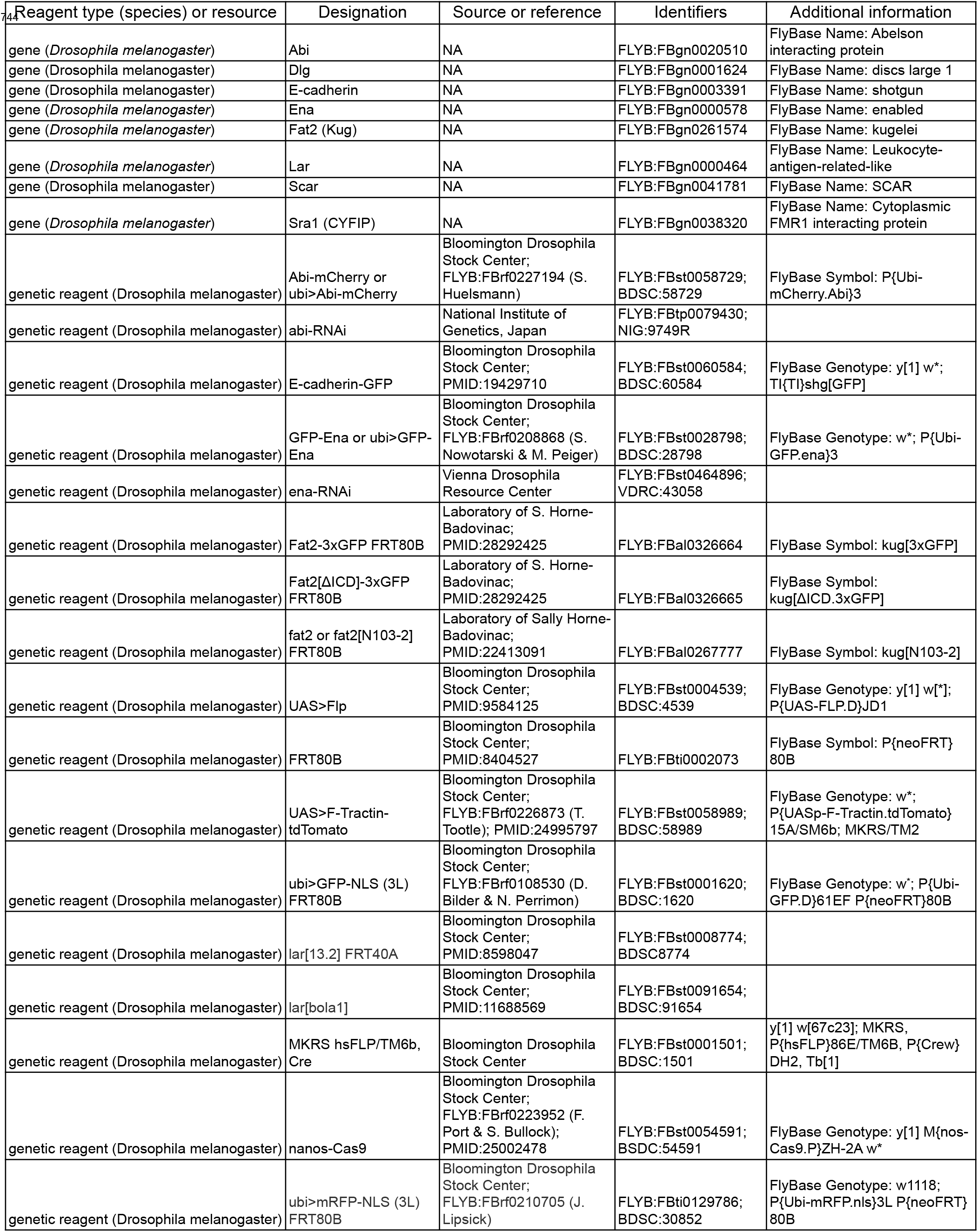
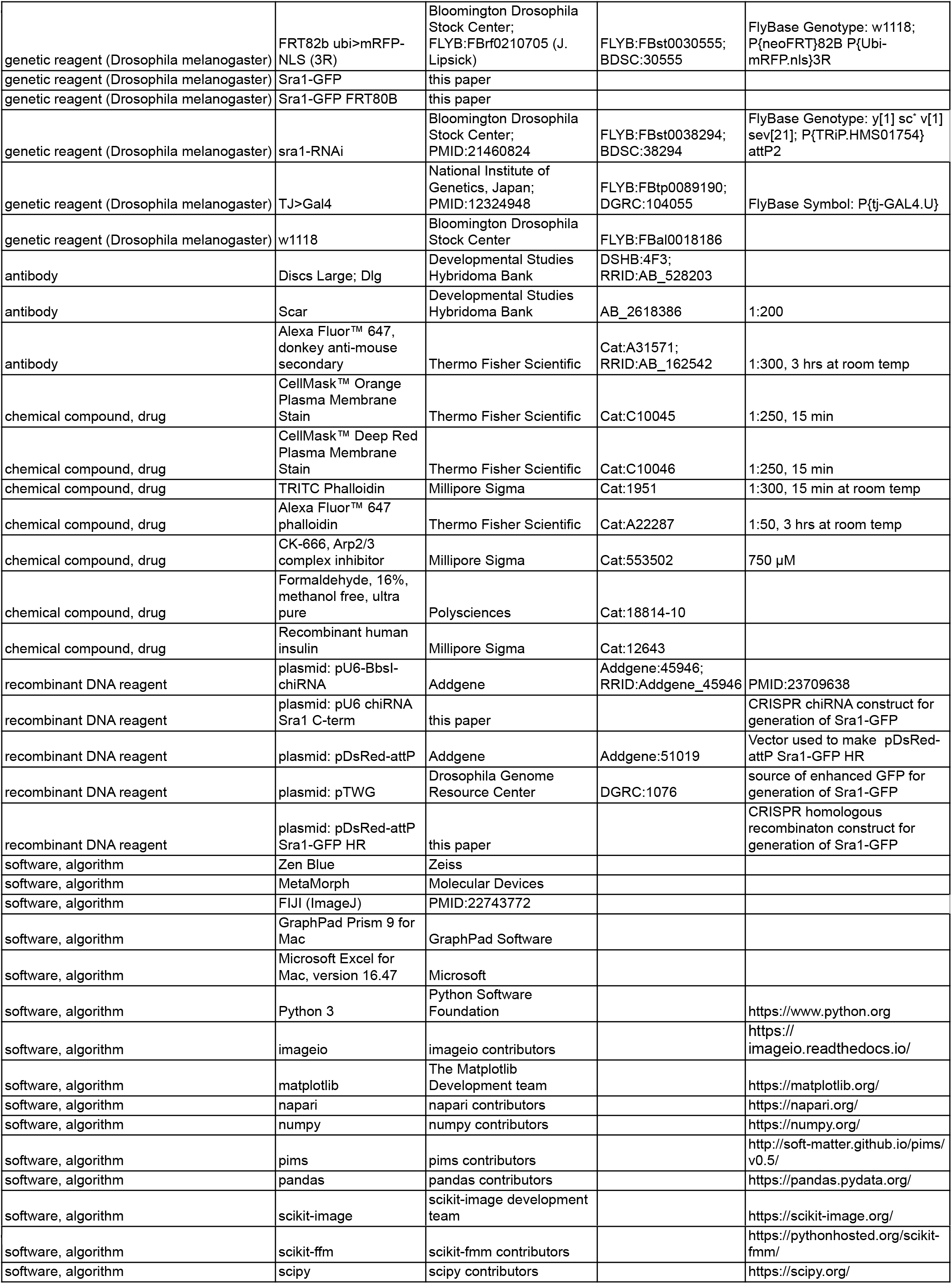
Key resources

**Supp. Table 2:** Genotypes of experimental females

| Figure | Panel | Name | Genotype |
| --- | --- | --- | --- |
| 2<br>747 | A-C | Control | w1118 |
|  |  | *fat2* | w;; *fat2*<sup>N103-2</sup> FRT80B |
|  |  | CK-666 | w1118 |
|  | D | Control | w1118 |
|  |  | *fat2* | w;; *fat2*<sup>N103-2</sup> FRT80B |
| 3 | B | Sra1-GFP mosaic | w; tj>Gal4<sup>DGRC:104055</sup>, UAS>Flp<sup>BDSC:4539/+</sup>; FRT82B Sra1-GFP/FRT82B ubi>mRFP-NLS<sup>BDSC:30555</sup> |
|  | C | *fat2* mosaic | w; tj>Gal4<sup>DGRC:104055</sup>, UAS>Flp<sup>BDSC:4539/+</sup>; *fat2*<sup>N103-2</sup> FRT80B/ubi>GFP-NLS FRT80B<sup>BDSC:1620</sup> |
|  | D-F | *fat2* mosaic + Sra1 | w; tj>Gal4<sup>DGRC:104055</sup>, UAS>Flp<sup>BDSC:4539/+</sup>; *fat2*<sup>N103-2</sup> FRT80B Sra1-GFP/ubi>mRFP-NLS FRT80B<sup>BDSC:30852</sup> |
| 4 | A | Sra1-GFP mosaic + *fat2* | w; tj>Gal4<sup>DGRC:104055</sup>, UAS>Flp<sup>BDSC:4539/+</sup>; *fat2*<sup>N103-2</sup> FRT80B FRT82B Sra1-GFP/*fat2*<sup>N103-2</sup> FRT80B FRT82B |
|  | B,D | Control | w;; Sra1-GFP/+ |
|  |  | *fat2* | w;; *fat2*<sup>N103-2</sup> FRT80B Sra1-GFP/*fat2*<sup>N103-2</sup> FRT80B |
|  | E | Control | w1118 |
|  |  | *fat2* | w; *fat2*<sup>N103-2</sup> FRT80B |
| 5 | A,E | Fat2 + Abi | w;; ubi>Abi-mCherry<sup>BDSC:58729</sup>, Fat2-3xGFP FRT80B/ Fat2-3xGFP FRT80B |
|  |  | Fat2<sup>ΔICD</sup> + Abi | w;; ubi>Abi-mCherry<sup>BDSC:58729</sup>, Fat2<sup>ΔICD</sup>-3xGFP FRT80B/Fat2-3xGFP FRT80B |
|  | B | Fat2 + Abi | w;; ubi>Abi-mCherry<sup>BDSC:58729</sup>, Fat2-3xGFP FRT80B/ Fat2-3xGFP FRT80B |
|  | C,D,F,G | Fat2 + Abi + F-actin | w;; ubi>Abi-mCherry<sup>BDSC:58729</sup>, Fat2-3xGFP FRT80B/ Fat2-3xGFP FRT80B |
| S1 | Top row | Protrusion in 1 direction | w1118 |
|  | Bottom row | Protrusion in both directions | w; *fat2*<sup>N103-2</sup> FRT80B |
| S2 | A | Control | w1118 |
|  |  | *fat2* | w;; *fat2*<sup>N103-2</sup> FRT80B |
|  |  | CK-666 | w1118 (treated with 750 μM CK-666) |
|  | B | Control | w1118 |
|  |  | *fat2* | w;; *fat2*<sup>N103-2</sup> FRT80B |
| S3 | A-C | Control | w; tj>Gal4<sup>DGRC:104055/+</sup> |
|  |  | *fat2* | w; tj>Gal4<sup>DGRC:104055/+</sup>; *fat2*<sup>N103-2</sup> FRT80B |
|  |  | *abi*-RNAi | w; tj>Gal4<sup>DGRC:104055/+</sup>; UAS>*abi*-RNAi<sup>NIG:9749R-3/+</sup> |
|  | D | Control | w; tj>Gal4<sup>DGRC:104055/UAS>F-Tractin-tdTomato</sup><sup>BDSC:58989</sup> |
|  |  | *fat2* | w; tj>Gal4<sup>DGRC:104055/UAS>F-Tractin-tdTomato</sup><sup>BDSC:58989</sup>; *fat2*<sup>N103-2</sup> FRT80B |
|  |  | *abi*-RNAi | w; tj>Gal4<sup>DGRC:104055/UAS>F-Tractin-tdTomato</sup><sup>BDSC:58989</sup>; UAS>*abi*-RNAi<sup>NIG:9749R-3/+</sup> |
|  | E,F | Control | w1118 |
|  |  | *fat2* | w;; *fat2*<sup>N103-2</sup> FRT80B |
| S4 | A,B | *fat2* mosaic | w; tj>Gal4<sup>DGRC:104055</sup>, UAS>Flp<sup>BDSC:4539/+</sup>; *fat2*<sup>N103-2</sup> FRT80B/ubi>GFP-NLS FRT80B<sup>BDSC:1620</sup> |
| S5 | A | Sra1-GFP | w;; Sra1-GFP |
|  |  | anti-SCAR | w1118 |
|  |  | ubi>Abi-mCherry | w;; ubi>Abi-mCherry<sup>BDSC:58729/+</sup> |
|  | B | Control | w; tj>Gal4<sup>DGRC:104055/+</sup> |
|  |  | *abi*-RNAi | w; tj>Gal4<sup>DGRC:104055/+</sup>; UAS>*abi*-RNAi<sup>NIG:9749R-3/+</sup> |
|  | C | Control | w1118 |
|  |  | Sra1-GFP x1 | w;; Sra1-GFP/+ |

Supp. Table 2: Genotypes of experimental females
|  |  |  |  |
| --- | --- | --- | --- |
| 748 |  | Sra1-GFP x2 | w;; Sra1-GFP |
|  |  | *sra1*-RNAi | w; tj>Gal4<sup>DGRC:104055/+</sup>; UAS>*sra1*-RNAi<sup>BDSC:38294/+</sup> |
|  | D | Control | w1118 |
|  |  | Sra1-GFP x1 | w;; Sra1-GFP/+ |
|  |  | Sra1-GFP x2 | w;; Sra1-GFP |
| S6 | A,C-E | Control | w;; Sra1-GFP/+ |
|  |  | *fat2* | w;; *fat2*<sup>N103-2</sup> FRT80B Sra1-GFP/*fat2*<sup>N103-2</sup> FRT80B |
| S7 | A | Control | w;; Sra1-GFP/+ |
|  |  | *fat2* | w;; *fat2*<sup>N103-2</sup> FRT80B Sra1-GFP/*fat2*<sup>N103-2</sup> FRT80B |
|  | B,C | *fat2* mosaic + Sra1 | w; tj>Gal4<sup>DGRC:104055</sup>, UAS>Flp<sup>BDSC:4539/+</sup>; *fat2*<sup>N103-2</sup> FRT80B Sra1-GFP/ubi>mRFP-NLS FRT80B<sup>BDSC:30852</sup> |
| S8 | A,B | Control Fat2 + Abi | w;; ubi>Abi-mCherry<sup>BDSC:58729</sup>, Fat2-3xGFP FRT80B/<br>Fat2-3xGFP FRT80B |
|  |  | *ena*-RNAi, Fat2 + Abi | w; tj>Gal4<sup>DGRC:104055</sup>/UAS>*ena*-RNAi<sup>VDRC:43058</sup>; ubi>Abi-mCherry<sup>BDSC:58729</sup>, Fat2-3xGFP FRT80B/<br>Fat2-3xGFP FRT80B |
|  |  | *lar* Fat2 + Abi | w; *lar*<sup>bola1</sup> BDSC:91654/*lar*<sup>13.2</sup> BDSC8774 FRT40A; ubi>Abi-mCherry<sup>BDSC:58729</sup>, Fat2-3xGFP FRT80B/<br>Fat2-3xGFP FRT80B |
|  |  | Control E-cadherin + Abi | w; Shg-GFP<sup>BDSC:60584/+</sup>; ubi>Abi-mCherry<sup>BDSC:58729/+</sup> |
|  | C-E | Ena + Abi + F-actin | w; ubi>GFP-Ena<sup>BDSC:28798</sup>/ubi>Abi-mCherry<sup>BDSC:58729</sup> |
|  | F | Control | w; tj>Gal4<sup>DGRC:104055/+</sup> |
|  |  | *ena*-RNAi | w; tj>Gal4<sup>DGRC:104055</sup>/UAS-*ena*-RNAi<sup>VDRC:43058</sup> |
|  |  | *abi*-RNAi | w; tj>Gal4<sup>DGRC:104055/+</sup>; UAS-*abi*-RNAi<sup>NIG:9749R-3/+</sup> |
|  | G | Control | w;; ubi>Abi-mCherry<sup>BDSC:58729</sup>, Fat2-3xGFP FRT80B/<br>Fat2-3xGFP FRT80B |
|  |  | *lar* | w; *lar*<sup>bola1</sup> BDSC:91654/*lar*<sup>13.2</sup> BDSC8774 FRT40A; ubi>Abi-mCherry<sup>BDSC:58729</sup>, Fat2-3xGFP FRT80B/<br>Fat2-3xGFP FRT80B |
| S9 |  | Fat2 + Abi | w;; ubi>Abi-mCherry<sup>BDSC:58729</sup>, Fat2-3xGFP FRT80B/<br>Fat2-3xGFP FRT80B |

**Supp. Table 3:**
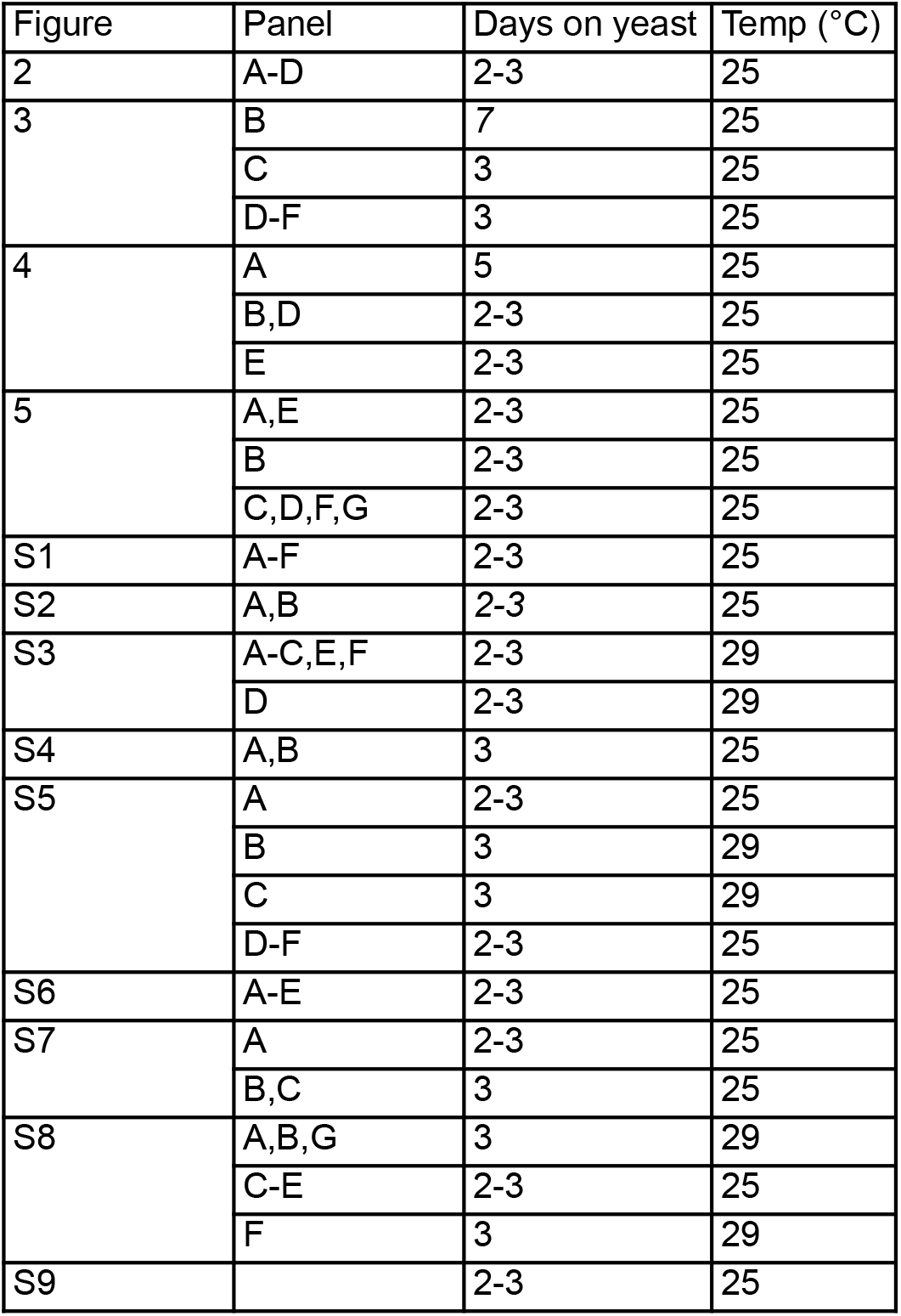
*Drosophila* culture conditions for experiments

